# Genotyping-by-sequencing reveals the effects of riverscape, climate and interspecific introgression on the genetic diversity and local adaptation of the endangered Mexican golden trout (*Oncorhynchus chrysogaster*)

**DOI:** 10.1101/500041

**Authors:** Marco A. Escalante, Charles Perrier, Francisco J. García-De León, Arturo Ruiz-Luna, Enrique Ortega-Abboud, Stéphanie Manel

## Abstract

How environmental and anthropogenic factors influence genetic variation and local adaptation is a central issue in evolutionary biology. The Mexican golden trout (*Oncorhynchus chrysogaster*), one of the southernmost native salmonid species in the world, is susceptible to climate change, habitat perturbations and the competition and hybridization with exotic rainbow trout (*O. mykiss*). The present study aimed for the first time to use genotyping-by-sequencing to explore the effect of genetic hybridization with *O. mykiss* and of riverscape and climatic variables on the genetic variation among *O. chrysogaster* populations. Genotyping-by-sequencing (GBS) was applied to generate 9767 single nucleotide polymorphisms (SNPs), genotyping 272 *O. chrysogaster* and *O. mykiss*. Population genomics analyses were combined with landscape ecology approaches into a riverine context (riverscape genetics). The clustering analyses detected seven different genetic groups (six for *O. chrysogater* and one for aquaculture *O. mykiss*) and a small amount of admixture between aquaculture and native trout with only two native genetic clusters showing exotic introgression. Latitude and precipitation of the driest month had a significant negative effect on genetic diversity and evidence of isolation by river resistance was detected, suggesting that the landscape heterogeneity was preventing trout dispersal, both for native and exotic individuals. Moreover, several outlier SNPs were identified as potentially implicated in local adaptation to local hydroclimatic variables. Overall, this study suggests that *O. chrysogater* may require conservation planning given i) exotic introgression from *O. mykiss* locally threatening *O. chrysogater* genetic integrity, and ii) putative local adaptation but low genetic diversity and hence probably reduced evolutionary potential especially in a climate change context.

## 1. INTRODUCTION

Natural and anthropogenic factors influence micro-evolutionary processes (i.e. migration, drift, and selection) generating spatial genetic patterns in wild populations. Disentangling the respective roles of environmental and anthropogenic variables in spatial genetic patterns has therefore become a central issue in evolutionary biology and conservation ecology (Baguette et al. 2013; Richardson et al. 2016).

Landscape genetics, and more specifically riverscape genetics (combination of population genetics with landscape ecology into a riverine context), can improve our understanding of the effects that riverine landscapes (riverscapes) and anthropogenic activities (e.g. aquaculture with exotic species) have on the microevolutionary processes determining the spatial genetic structure and genetic diversity of populations, particularly in lotic environments (Manel and Holderegger 2013; Davis et al. 2018; Grummer et al. 2019). Microevolutionary processes are influenced by different alterations such as landscape fragmentation and water flow changes, which usually reduce habitat size, split environments and create barriers that isolate fish populations (Morita and Yamamoto 2002; Le Pichon et al. 2006). Gene flow and selection act on different ways (i.e. among basins, and between distant or nearby sites within basins). Therefore, it is necessary to account for those differences to analyse riverine environments, particularly in complex hydrological networks where the physical landscape structure, along with the reduced pathways, determines patterns of genetic variation (Chaput-Bardy et al. 2008; Kanno et al. 2011).

Both neutral and adaptive components of genetic diversity are the raw materials of evolution and should be considered in conservation studies (Kokko et al. 2017; Tikochinski et al. 2018). However, most of the already published riverscape genetics studies used only a small number of neutral molecular markers. Lately, some studies started to include large Next Generation Sequencing (NGS) datasets to assess gene-environment interactions (Bourret et al. 2013; Hecht et al. 2015; Brauer et al. 2016, 2018; Hand et al. 2016; Amish et al. 2019). The use of NGS datasets might be helpful for the genetic monitoring of endangered populations, as they provide information of local adaptation processes that is valuable for recovery programmes which could be apply in riverine environments (Faulks et al. 2017; Kleinman-Ruiz et al. 2017; Hunter et al. 2018; Fan et al. 2018).

For riverine species the introduction of exotic species for recreational and aquaculture purposes, might have a detrimental effect on the native populations due to predation, competition, the introduction of diseases and genetic introgression (Perrier et al. 2011; Penaluna et al. 2016). When the exotic species is phylogenetically close to the native species it can cause genetic introgression, which in turn can have extremely harmful effects such as homogenizing biodiversity (Allendorf et al. 2001). Moreover, the modification of the native genetic pool might result in the loss of selective values (fitness) affecting survival skills in changing environments (Milián-García et al. 2015). Recent climatic variations have been associated with the spread of genetic introgression in riverine habitats, leading into irreversible evolutionary consequences for endangered species. The negative impact of genetic introgression in reverine habitats have been widely documented in salmonids due to the extensive use of exotic species for aquaculture (Perrier et al. 2013; Muhlfeld et al. 2017).

Climate conditions such as temperature and precipitation might be drivers of adaptive processes in different salmonid species because they are associated with migration, reproduction, and mortality. In this sense, upstream salmonids are extremely vulnerable to the detrimental effects arising from climate change, as they have a much smaller habitat than marine and terrestrial environments (Parmesan 2006; Muhlfeld et al. 2017); but also, because of their specific habitat requirements (Ruiz-Luna et al. 2017).

Such is the case of the Mexican golden trout (MGT, *Oncorhynchus chrysogaster*; Needham and Gard, 1964), one of the southernmost native salmonid species, inhabiting three different basins (Río Fuerte, Río Sinaloa, and Río Culiacán) in the highest parts of the Sierra Madre Occidental (SMO) mountains in north-western Mexico (Hendrickson et al. 2002). The most broadly accepted hypothesis to explain the origin of this species is the colonization process which occurred at the end of the Pleistocene (~ 12 000 years ago), when a group of steelhead trout migrated from the Gulf of California to the upper parts of SMO rivers (Behnke et al. 2002). Subsequently, these trout were isolated in different streams and basins experiencing genetic drift, reduced gene flow, and potentially specific selective pressure (García-De León et al. 2020). Like all salmonids, this species is susceptible to climate change, habitat perturbations, and the introduction of non-native species (i.e. rainbow trout for aquaculture purposes; Hendrickson et al. 2006). *O. chrysogaster* is an endangered species listed in the Mexican official norm of threatened species (NOM 059) and the International Union for Conservation of Nature (IUCN) red list. Many aspects of *O. chrysogaster*’s biology including its foraging habits, homing and reproductive behaviour, and dispersal are still unknown (but see Ruiz-Luna and García-De León, 2016). Biological studies of this species have mainly been limited due to the difficulty accessing its habitat because of both the lack of road infrastructure, and the prevalence of drug trafficking. Existing genetic studies of this species have suggested that *O. chrysogaster* is genetically structured by hydrological basins (e.g.; Camarena-Rosales et al. 2008; Escalante et al. 2014) and a recent works described up to four distinct genetic clusters (Escalante et al. 2016; García-De León et al. 2020). However, these studies were conducted with neutral markers, a small number of sample sites, and sample designs that were not adapted to account for riverscape complexity. This means that microevolutionary processes and their association with the adjacent riverscape are not yet properly understood, and consequently there is a lack of management plans to protect this endangered species.

Some studies for this species have revealed a genetic substructure and introgression processes in locations near aquaculture facilities (Escalante et al. 2014; Abadía-Cardoso et al. 2015). Moreover, a recent simulation study of *O. chrysogaster* suggested that the riverscape acts as a barrier against exotic introgression and that riverscape resistance (the combined effect of riverine distance, topographic slope, altitude gradient, stream order changes and temperature increase) is the main factor fragmenting populations within basins (Escalante et al. 2018). Strong spatial genetic structure and local adaptation processes in *O. chrysogaster* arising from riverscape factors are therefore to be expected, with the consequence that this complex riverscape will prevent introgression processes in populations not in proximity to aquaculture farms.

This study focuses on the effect of anthropogenic factors as well as riverscape and climatic variables on the genetic diversity, population structure and local adaptation of the endangered endemic salmonid *O. chrysogaster* in order to obtain technical data for developing management and conservation strategies. Specifically, it addresses the following questions: 1) What is the impact of *O. mykiss* aquaculture escapes on the genetic structure of the endemic *O. chrysogaster*? 2) Which riverscape features (i.e. hydroclimatic and topographic variables) affect the extent and distribution of genetic diversity? 3) Are there genomic footprints of local adaptation to heterogeneous hydroclimatic features? In order to answer these questions, this study explored the genetic structure of *O. chrysogaster* populations and aquaculture trout (237 wild trout, 7 aquaculture *O. chrysogaster*, 24 aquaculture *O. mykiss*, and 4 lab hybrids), collected from 28 sites in three basins in north-western Mexico or provided by federal agencies and genotyped for 9767 Single Nucleotide Polymorphisms (SNPs). The results were discussed in the context of conservation strategies for endangered riverine species, considering both ecological and evolutionary criteria.

## 2. MATERIALS AND METHODS

### 2.1. Study system and data collection

The survey was conducted in the Río Fuerte, Río Sinaloa, and Río Culiacán basins, at altitudes ranging from 1 965 to 2 730 m (Fig. 1). The study area covers approximately 27 500 km^2^ with an estimated length of the river network of 12,700 km. This zone has a complex topography and hydrology with sharp slopes and stream orders mostly corresponding to 1, 2 and 3. The annual mean temperature and annual mean precipitation for the zone are 14.4 °C and 911 mm respectively (Escalante et al. 2018). The heterogeneous riverscape of this area offers a diversity of terrestrial and aquatic habitats providing high endemism and biodiversity, with *O. chrysogaster* as one of the main aquatic predators (Hendrickson et al. 2006). For these river basins the dams are present downstream at altitudes of less than 500 meters, and at altitudes above 1000 meters there is no evidence of other anthropogenic undertakings, except for some rural activities (e.g. aquaculture and artisanal agriculture; author’s field observations). However, the introduction of rainbow trout for aquaculture purposes has been reported since the 1860s, and was strongly supported by federal agencies during the 1980s and 1990s (Escalante et al. 2014).

**Fig. 1.**
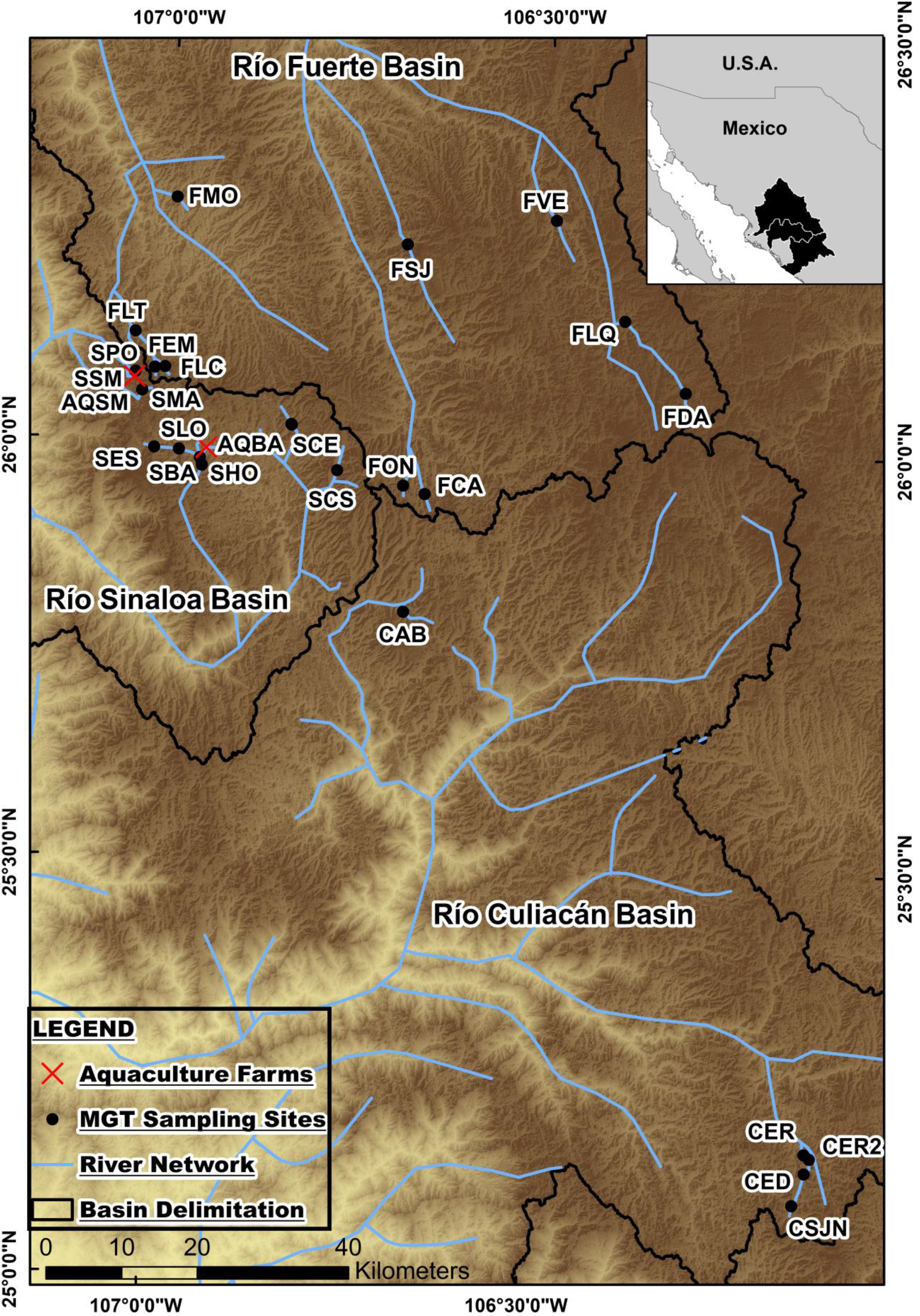
Study area. Mexican golden trout (MGT) and cultured rainbow trout (Aquaculture farms) sample sites in the Río Fuerte, Río Sinaloa and Río Culiacán basins. See table 1 for the explanation of the sample site codes

Six variables characterizing the riverscape and with a strong influence on *O. chrysogaster* occurrence and genetic divergence were obtained from a previous study (Escalante et al. 2018): precipitation of the driest month, temperature of the warmest quarter, river length (distance from the lowermost to the uppermost part of the river above 1 500 m), slope, altitude, and stream order. They were generated and/or processed from data available on the worldclim (http://www.worldclim.org/) and the Japanese Space System (http://www.jspacesystems.or.jp/ersdac/GDEM/E/4.html) websites with 1 km and 30 meter resolution, respectively. Worldclim variables were geometrically corrected to 30 meter resolution (for further details see Escalante et al. 2018). The effect of latitude and longitude, recorded using GPS, was also tested.

Wild trout were sampled by electrofishing at 25 sample sites in the Río Fuerte (10 sites), Río Sinaloa (10 sites), and Río Culiacán (5 sites) basins during the winter and spring seasons of 2013, 2014, and 2015 (collection permit numbers SGPA/DGVS/02485/13 and SGPA/DGVS/05052/15). The sampling was conducted from both the core and edges of the *O. chrysogaster* ecological niche defined in a previous study (Escalante et al. 2018), and at multiple locations with high environmental heterogeneity (topographical and hydrological divergence) avoiding spatial autocorrelation (Fig. 1). Rainbow trout samples were obtained from two aquaculture farms in the Río Sinaloa basin. Additionally, seven farmed *O. chrysogaster* samples and four lab hybrids from *O. chrysogaster* and *O. mykiss* from Guachochi Aquculture Center in Maderas Mexico donated by the Mexican Institute of Fisheries (INAPESCA), were included in the study for further genotyping. Small portions of tissue (either fin or muscle) were clipped from these samples and preserved in 95% ethanol for later analysis.

### 2.2. Genotyping by sequencing

Genomic DNA was extracted for each individual following the Qiagen DNeasy Blood & Tissue Kit protocol (Qiagen, Hilden, Germany, http://www1.qiagen.com). DNA quality was checked using agarose gel electrophoresis and quantified using a NanoDrop spectrophotometer (Thermo Scientific) and the Quant-iT PicoGreen dsDNA Assay Kit (Invitrogen). DNA libraries were then generated with GBS methods (Elshire et al. 2011) at Cornell University in Ithaca, New York, using the ECOT221 enzyme. Finally, single-read 100-bp sequencing was performed on an Illumina HiSeq2500.

The quality of the raw sequences was controlled via investigating per base averaged quality, presence of adapters and over-represented sequences were detected using FastQC (Andrews 2010). Reads were processed with Cutadapt (Martin 2011) to remove potential fragments of Illumina adapters, allowing a 10% mismatch rate in the adapter sequence. During this process, reads were trimmed to 70 bp. The bioinformatics software/pipeline Stacks 1.32 (Catchen et al. 2011; Catchen 2013) was hence used to demultiplex reads, identify restriction site-associated DNA (RAD) loci and call SNPs. Reads were demultiplexed and trimmed to 64 bp using the process_radtags module, with which, one potential mismatch in the barcode sequence was allowed. The ustacks module was used, with a minimum stack depth of 4x and a maximum distance allowed between stacks of 4 (6 for secondary reads). The catalogue of loci was built using the cstacks module with n = 4. With the sstacks module, samples were matched against the catalogue of loci. Finally, individuals were genotyped using the population module, with at least 70% of individuals being genotyped, and a minimum read depth of 5x for each locus and individual. Genotypes were exported in VCF format for further filtering. The SNP dataset was filtered using VCFtools (Danecek et al. 2011), achieving a minimum average read depth ranging from 8x to 40x across genotypes and a minor allele frequency of 1%, to limit any potential low quality data or paralogous loci. Based on three populations with large sampling sizes that were exempt of stocking (FVE, FLQ and SBA) a blacklist of loci deviating from Hardy-Weinberg equilibrium (HWE, p-value ≤ 0.05) was constituted. Thus, loci deviating from HWE in at least one of the three populations considered were removed.

Subsequently, five datasets were created including all loci after SNP calling at four different spatial scales: Dataset A for population genetics analyses including all the genotyped individuals; Dataset B for gene-environment associations (landscape genomics analyses) across the entire study area without significantly introgressed native trout (individuals with ancestry coefficients ≥ 20% from aquaculture were removed); Dataset C for landscape genetics analyses with spatially continuous native populations (removing the isolated populations in the south of Río Culiacán to prevent biases in the landscape genetics analyses) without significant introgression; Dataset D for landscape genetics analyses of native trout without significant introgression in Río Fuerte (removing sampling sites isolated by long riverine distance at the east of Río Fuerte); and Dataset E for landscape genetics analyses of native trout without significant introgression in Río Sinaloa. Further information about the datasets is included in Online Resource 1.

### 2.3. Population genomics analyses

The genetic diversity of trout (Dataset A) was estimated from expected heterozygosity (*H_E_*), and observed heterozygosity (*H_O_*) using the adegenet R package (Jombart, 2008). Effective population size (*N_E_*) was calculated by applying a molecular co-ancestry method (Nomura 2008) implemented in NeEstimator (Do et al. 2014).

The genetic differentiation between all sample sites (Dataset A) was assessed considering three different approaches. Firstly, pairwise genetic differentiation coefficients (*F_ST_*) were calculated between all sample sites in Genodive 3.0 (Meirmans and Van Tienderen 2004); this method uses an analysis of molecular variance, performed between each population pair. Secondly, a phenogram was built in adegenet from all individuals using the neighbour-joining algorithm (Saitou and Nei 1987) based on Nei’s genetic distance (Tamura and Nei 1993). Bootstrap confidence intervals were estimated from 10 000 permutations.

Thirdly, an unsupervised Bayesian clustering approach implemented in the package fastStructure (Raj et al. 2014) was used to assess the genetic structure of *O. chrysogaster* and genetic admixture with aquaculture trout. This approach infers population genetic structure for a large number of SNP datasets without assuming predefined populations. Thus, fastStructure was run using Dataset A including all SNPs from all the genotyped individuals and Dataset B including all SNPs from individuals without significant exotic introgression. Based on an approximation of the number of sample sites (total number of sampling sites + 1), *K* = 30 was considered the maximum value to avoid underestimating the number of clusters.

### 2.4. Riverscape genetics analyses

Only Dataset C was considered when exploring the riverscape’s influence on genetic diversity, due to the reduced number of sampling sites for the datasets at intra-basin scale (i.e. datasets D and E). Initially, the collinearity between seven predictors was tested using the R package corrplot (Taiyun Wei 2017): latitude, longitude, precipitation of the driest month, temperature of the warmest quarter, river length, altitude, and stream order. Afterwards, to test the effect of both climate and geographical distance, only latitude and precipitation of the driest month were retained because of their low collinearity (−0.48). Finally, the effect of latitude and precipitation of the driest month on expected heterozygosity was tested by a generalized linear model using the GLM function in R (Team R. Core 2018).

In order to examine the effect of riverscape on genetic differentiation, a resistance surface was firstly derived from four riverscape features using ArcGIS v10.2 (ESRI, 2013). We chose to adopt a resistance surface approach accounting for the combined effect of riverine distance, slope, altitude gradient, stream order changes and temperature increase as it was suggested to be a determinant factor driving *O. chrysogaster* genetic structure in a recently published simulation study (Escalante et al. 2018). A single point based environmental analysis is not able to quantify such physical corridors or boundaries to gene flow (Cushman and Landguth 2010; Landguth et al. 2012, 2016; Milanesi et al. 2017; Grummer et al. 2019). This surface was defined using the temperature of the warmest quarter, slope, stream order, and altitude raster files. Values were assigned to the pixels at each raster representing the extent to which movement is obstructed. The same criteria as those reported by Escalante et al. (2018) were considered, as the species and the study area fit with the current study. Then, resistance values from 0 to 10 were assigned to the raster pixels (30 m resolution) for each variable independently (see Online Resource 2). A riverscape resistance raster was generated by averaging the resistance values for the four environmental variables at each pixel. For subsequent analyses in the gdistance package (van Etten, 2012), resistance values were rescaled from 1 to 2, with 1 representing an absence of riverscape resistance and 2 maximum resistance. Further information about the parameterisation is included in Online Resource 2.

The gdistance R package (van Etten 2012) was then used to calculate linear distance and riverscape resistance matrices among sample sites within basins. This method simulates potential species movement in a spatially structured landscape, linking different dispersal functions and connectivity thresholds using Djikstra’s shortest path algorithm (Dijkstra 1959). Two matrices were generated under the hypothesis of isolation by Riverscape Resistance (RR) at intra-basin scale: RR for Río Fuerte populations (Matrix I) and RR for Río Sinaloa populations (Matrix II). Additionally, two linear distance matrices were generated for the Río Fuerte populations (Matrix III) and Río Sinaloa populations (Matrix IV) in order to control differences in distance in subsequent analyses (for further details, see Online Resource 2). To define the influence of isolation by riverscape resistance on genetic distances, Partial Mantel tests were applied using the PASSaGE package (Rosenberg and Anderson 2011). Partial Mantel tests were performed between the regression of *F_ST_*/(1- *F_ST_*) of Dataset D (Río Fuerte) and Dataset E (Río Sinaloa) and corresponding riverscape resistance matrices at intra-basin scale (either Matrix I, either Matrix II), using geographical linear distance as constant matrices (either Matrix III, either Matrix IV) allowing the control for differences in distance (Online Resource 2). All tests were performed on under 10 000 permutations and assuming no correlation.

### 2.5. Detection of SNPs under divergent selection

Dataset B was analysed to detect *O. chrysogaster* SNPs potentially under selection using three different software programs and two different approaches: a population (i.e. sampled sites) outlier detection approach (1) and association tests between genotypes and continuous climatic variables (i.e. riverscape adaptive genomics) (2).

Firstly, the PCAdapt R package (Luu et al. 2016) was used to detect SNPs potentially under divergent selection with approach 1. This method combines principal component analysis and Mahalanobis distances and assumes that molecular markers excessively associated with population structure are candidates for local adaptation. Based on the vector of z-scores, loci which do not follow the distribution of the main cluster of points are considered outliers. The analysis was run with a threshold of 10% and K=6 based on fastStructure results (observed spatial genetic structure).

For approach 2, two gene-environment association software programs were used to test the effect of temperature of the warmest quarter and precipitation of the driest month. These variables had previously been suggested as significant adaptation drivers in salmonids (Hecht et al. 2015; Hand et al. 2016). Using mixed models, both methods detect outlier loci through allele frequencies exhibiting strong statistical correlations with environmental variables. Initially, Bayenv2 was applied (Günther and Coop 2013), using an average of five independent runs (100 000 iterations). Also, the latent factor mixed models (LFMM) algorithm implemented in the R package LEA (Frichot et al. 2013; Frichot and François 2015) was run with five repetitions, 10 000 cycles, 5 000 burn-in, and K=6 (based on fastStructure outputs of the spatial genetic structure) as a random factor in the regression analysis. For both LFFM and Bayenv2, a threshold of 1% of total SNPs was defined to select the outlying SNPs with the highest posterior probabilities.

A gene ontology analysis was conducted on the nucleotide sequences (64 bp) containing all SNPs obtained from *O. chrysogaster*, using Dataset B. This analysis is based on a BLAST query (blastn) of the sequences against peptides from the rainbow trout genome database (Berthelot et al. 2014). Functional categorization by gene ontology terms (GO; http://www.geneontology.org) was carried out using Blast2GO software (version 4.1, http://www.blast2go.com/). Subsequently, protein annotations were filtered, retaining those from loci detected to be under divergent selection with an e-value cutoff of ≤ 10^−6^.

## 3. RESULTS

### 3.1. Genotyping by sequencing

The total number of raw sequences obtained was 722 836 026 with an average of 1 300 000 reads per individual. A genotyping rate of 82% was obtained for the total number of individuals sampled per population. After SNP calling, a total of 270 individuals (Table 1) and 9767 SNPs were retained for subsequent analysis.

**TABLE 1.** Genetic diversity values for Mexican golden trout, aquaculture rainbow trout and lab hybrids.

| River Basin | CODE | Latitude | Longitude | Location | Description | N<sup>a</sup> | H<sub>O</sub><sup>b</sup> | H<sub>E</sub><sup>c</sup> | N<sub>E</sub><sup>d</sup> | 95%<br>CI N<sub>E</sub><sup>e</sup> |
| --- | --- | --- | --- | --- | --- | --- | --- | --- | --- | --- |
| Río Fuerte | FDA | 26.07 | -106.31 | Arroyo del Agua | Wild trout | 10 | 0.03 | 0.04 | 44.2 | 38.3–51.9 |
|  | FEM | 26.09 | -107.02 | Arroyo El Manzano | Wild trout | 8 | 0.06 | 0.06 | 35.3 | 32.4–38.7 |
|  | FLC | 26.09 | -107.00 | Arroyo Las Cuevas | Wild trout | 10 | 0.06 | 0.07 | 33.7 | 32.2–35.3 |
|  | FLQ | 26.16 | -106.40 | Arroyo La Quebrada | Wild trout | 12 | 0.04 | 0.05 | 116.8 | 110.3–139.6 |
|  | FLT | 26.13 | -107.04 | Arroyo Las Truchas | Wild trout | 8 | 0.04 | 0.04 | - | Infinite |
|  | FMO | 26.29 | -106.99 | Arroyo Momorita | Wild trout | 5 | 0.02 | 0.02 | - | Infinite |
|  | FSJ | 26.24 | -106.69 | Arroyo San José | Wild trout | 12 | 0.07 | 0.07 | 14.7 | 14.4–15 |
|  | FCA | 25.94 | -106.65 | Arroyo Calera | Wild trout | 9 | 0.14 | 0.13 | 8.2 | 8.1–8.2 |
|  | FON | 25.95 | -106.68 | Arroyo La Onza | Wild trout | 6 | 0.14 | 0.12 | - | Infinite |
|  | FVE | 26.28 | -106.49 | Río Verde | Wild trout | 12 | 0.05 | 0.05 | 11 | 10.8–11.2 |
| Totals and means |  | - | - | - | - | 92 | 0.065 | 0.065 | 37.7 |  |
| Río Sinaloa | SBA | 25.98 | -106.95 | Arroyo Baluarte | Wild trout | 9 | 0.07 | 0.09 | 145.7 | 114.8–198.7 |
|  | SES | 25.99 | -107.01 | Arroyo El Soldado | Wild trout | 9 | 0.08 | 0.08 | 5.9 | 5.8–6 |
|  | SHO | 25.97 | -106.95 | Arroyo Hondo | Wild trout | 11 | 0.09 | 0.11 | 2.4 | 2.3–2.4 |
|  | SLO | 25.99 | -106.98 | Arroyo La Osera | Wild trout | 10 | 0.08 | 0.09 | 1.5 | 1.5–1.5 |
|  | SMA | 26.06 | -107.03 | Arroyo Macheras | Wild trout | 10 | 0.02 | 0.02 | 0.4 | 0.4–0.4 |
|  | SPO | 26.08 | -107.04 | Arroyo Potrero | Wild trout | 11 | 0.02 | 0.02 | - | Infinite |
|  | SSM | 26.07 | -107.04 | Arroyo San Miguel | Trout in aquaculture proximities | 4 | 0.17 | 0.15 | - | Infinite |
|  | SCS | 25.97 | -106.77 | Arroyo Cerro Solo | Wild trout | 9 | 0.11 | 0.15 | 1.4 | 1.4–1.4 |
|  | SCE | 26.02 | -106.83 | Arroyo Cebollín | Wild trout | 7 | 0.09 | 0.12 | 1.8 | 1.8–1.8 |
|  | SPE | 26.02 | -106.83 | Arroyo Pericos | Wild trout | 7 | 0.12 | 0.12 | 7.5 | 7.3–7.6 |
|  | **Totals and means** | - | - | - | - | **87** | **0.085** | **0.095** | **20.83** |  |
| **Río Culiacán** | CED | 25.14 | -106.13 | Arroyo El Desecho | Wild trout | 12 | 0.009 | 0.01 | - | Infinite |
|  | CER | 25.16 | -106.12 | Arroyo El Río 1 | Wild trout | 12 | 0.008 | 0.01 | - | Infinite |
|  | CER2 | 25.17 | -106.13 | Arroyo El Río 2 | Wild trout | 12 | 0.009 | 0.01 | - | Infinite |
|  | CSJN | 25.10 | -106.14 | Arroyo San Juan del Negro | Wild trout | 12 | 0.05 | 0.04 | 8 | 7.9-8.2 |
|  | CAB | 25.80 | -106.68 | Arroyo Agua Blanca | Wild trout | 10 | 0.11 | 0.1 | 42.7 | 41.4-44 |
|  | **Totals and means** | - | - | - | - | **58** | **0.038** | **0.034** | **25.35** |  |
| **Lab Trout** | H |  |  | INAPESCA Cultive Center | Mexican golden trout and aquaculture rainbow trout lab hybrids | 4 | 0.22 | 0.15 | - | Infinite |
|  | REP |  |  | INAPESCA Cultive Center | Domestic Mexican golden trout | 7 | 0.08 | 0.08 | - | Infinite |
|  | **Totals and means** | - | - | - | - | **11** | **0.15** | **0.12** |  |  |
| **Aquaculture Farms** | AQEB | 25.99 | -106.95 | El Barro Aquaculture Farm | Aquaculture rainbow trout | 14 | 0.15 | 0.16 | 19.1 | 18.9-19.2 |
|  | AQSM | 26.07 | -107.04 | San Miguel Aquaculture Farm | Aquaculture rainbow trout | 10 | 0.13 | 0.15 | 22.1 | 21.7-22.5 |
|  | **Totals and means** | - | - | - | - | **24** | **0.14** | **0.16** | **20.6** |  |

*a*-Number of genotyped samples (*N*), *b*-expected heterozygosity ( $H_E$ ), *c*-observed heterozygosity ( $H_O$ ), *d*-effective population size ( $N_E$ ) and *e*-95% confidence intervals for effective population size (95% CI $N_E$ ).

### 3.2. Population genomics analyses

The highest *H*_*O*_ values were observed in *O. chrysogaster* and rainbow trout lab hybrids (0.22) whereas wild *O. chrysogaster* collected at CED, CER, and CER2 in Río Culiacán obtained the lowest scores (0.01). For *H_E_*, the highest scores (0.16) were found in aquaculture trout collected at AQEB in Río Sinaloa, while the lowest values (0.01) were observed at CED, CER, and CER2 in Río Culiacán (Table 1). Focusing on wild *O. chrysogaster* only, the highest heterozygosity (both *H_E_* and *H_O_*) was found in the centre of the study area where the three basins meet (Fig. 2). *N_E_* were successfully estimated for 19 sample sites; the highest value (145.7) was found at SBA and the lowest (0.4) at SMA, both in Río Sinaloa (Table 1).

**Fig. 2.**
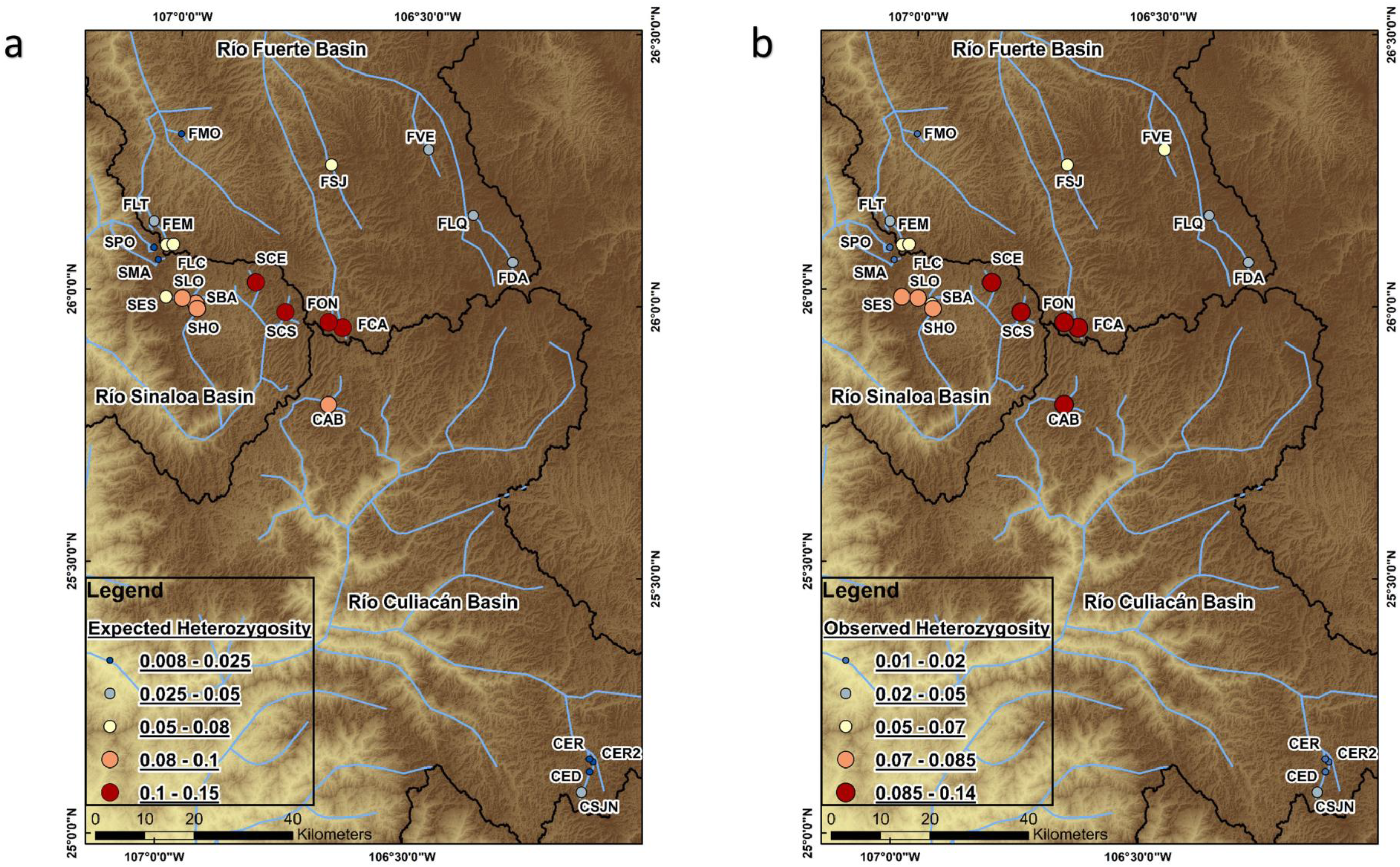
Geographical distribution of genetic diversity values for native *O. chrysogaster*. a) Expected heterozygosity; b) Observed heterozygosity. See table 1 for the explanation of the sample site codes

At the sampling sites, pairwise *F_ST_* was highest (0.964) at the SPO and CED sample sites, in two different basins (Río Culiacán and Río Sinaloa), while the lowest (0.008) was between CER and CER2 (both in Río Culiacán). In terms of the basins, the highest *F_ST_* averages (0.83) were observed between Río Fuerte and Río Culiacán (for further information, see Online Resource 3). The high *F_ST_* values in some populations reflect high spatial genetic structure and are normally observed in trout inhabiting heterogeneous landscapes with rugged topography and complex hydrology (e.g. Abadía-Cardoso et al. 2015) or in small isolated populations (Perrier et al. 2017).

The Nei phenogram detected 27 well-supported clades (>97 % of bootstrapping value), all made up of geographically close sampling sites. Interestingly, the aquaculture trout (AQEB and AQSM) showed greater genetic similarity (given its lower branch length) with the lab hybrids (H, clearly a heterogeneous group, as it is known that hybrids have been artificially obtained). Furthermore, the trout collected in SSM falls within the aquaculture clade, suggesting that they are rainbow trout escaped from the farming facilities (see below, Fig. 3).

**Fig. 3.**
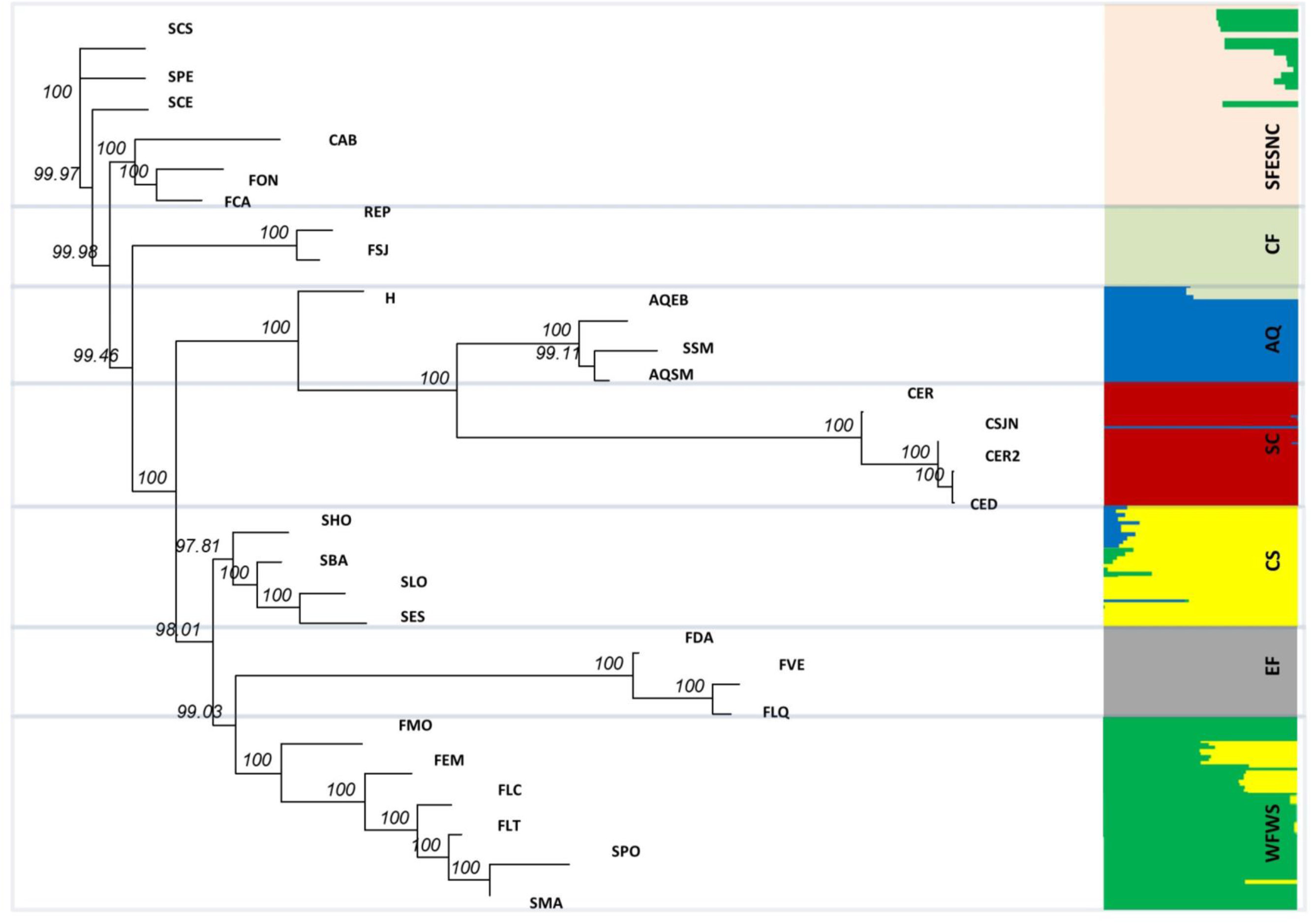
Genetic structure of *O. chrysogaster* and aquaculture rainbow trout defined by Nei distances (Tamura and Nei, 1993) in adegenet and a Bayesian assignment test in fastStructure. (a) Nei phenogram, the bootstrapping values for each clade are represented by numbers, see table 1 for the explanation of the sampling site codes. (b) Bayesian assignment test barplot, each color represents a different genetic cluster and the genome of each individual is represented by a horizontal line: Eastern Fuerte (EF); Central Fuerte (CF); Southern Fuerte, Eastern Sinaloa and Northern Culiacán (SFESNC); Western Fuerte and Western Sinaloa (WFWS); Central Sinaloa (CS); Southern Culiacán (SC); Aquaculture (AQ).

The unsupervised Bayesian clustering approach (fastStructure) identified six distinct wild genetic clusters defined by geography, and one aquaculture cluster: I. Eastern Fuerte; II. Central Fuerte; III. Southern Fuerte, Eastern Sinaloa, and Northern Culiacán; IV. Western Fuerte and Western Sinaloa; V. Central Sinaloa; VI. Southern Culiacán; and VII. Aquaculture (Fig. 3b and 4). Trout from Western Fuerte collected at FEM, FLC, and FLT showed admixture with trout from Central Sinaloa. Trout from Central Sinaloa collected at SBA showed genetic admixture with trout from Western Fuerte, while the trout collected at SHO and one individual from SLO showed admixture with aquaculture trout. Trout from Eastern Sinaloa collected at SCE, SCS, and SPE exhibited genetic admixture with Western Fuerte. Some trout from Southern Culiacán collected at CSJN exhibited genetic admixture with aquaculture trout, and one individual belonged entirely to the aquaculture cluster. Trout collected at SSM near the San Miguel aquaculture farm belonged to the aquaculture cluster and were identified as aquaculture trout, perhaps as a product of aquaculture escapes. Finally, for the lab hybrids, half of their genome belonged to native *O. chrysogaster* from the Central Fuerte cluster (parental population), and the other half to the aquaculture cluster.A comparison of different K values shows a similar genetic structure (across K=6, K=7, and K=8) with slight differences for K=6, where trout from FSJ (Central Fuerte in K=7 and 8) were clustered with trout from FCA, FON, CAB, SCE, SPE, and SCS (Online Resource 4). In general, the genetic admixture between native and farmed trout was very low. Moreover, for two genetic clusters (i.e. Southern Fuerte, Eastern Sinaloa, and Northern Culiacán; and Western Fuerte and Western Sinaloa; Fig. 3b and 4c) the observed spatial genetic structure does not follow a basin delineating, suggesting human translocations of native trout or potential connectivity among basins during flood events, or both. Similar results were observed while analysing Dataset B, obtaining six native genetic clusters with MGT individuals belonging to the same genetic clusters described above (Online Resource 5). Interestingly, analysing the dataset without outliers (with only neutral SNPs) found the same genetic structure (results not shown).

**Fig. 4.**
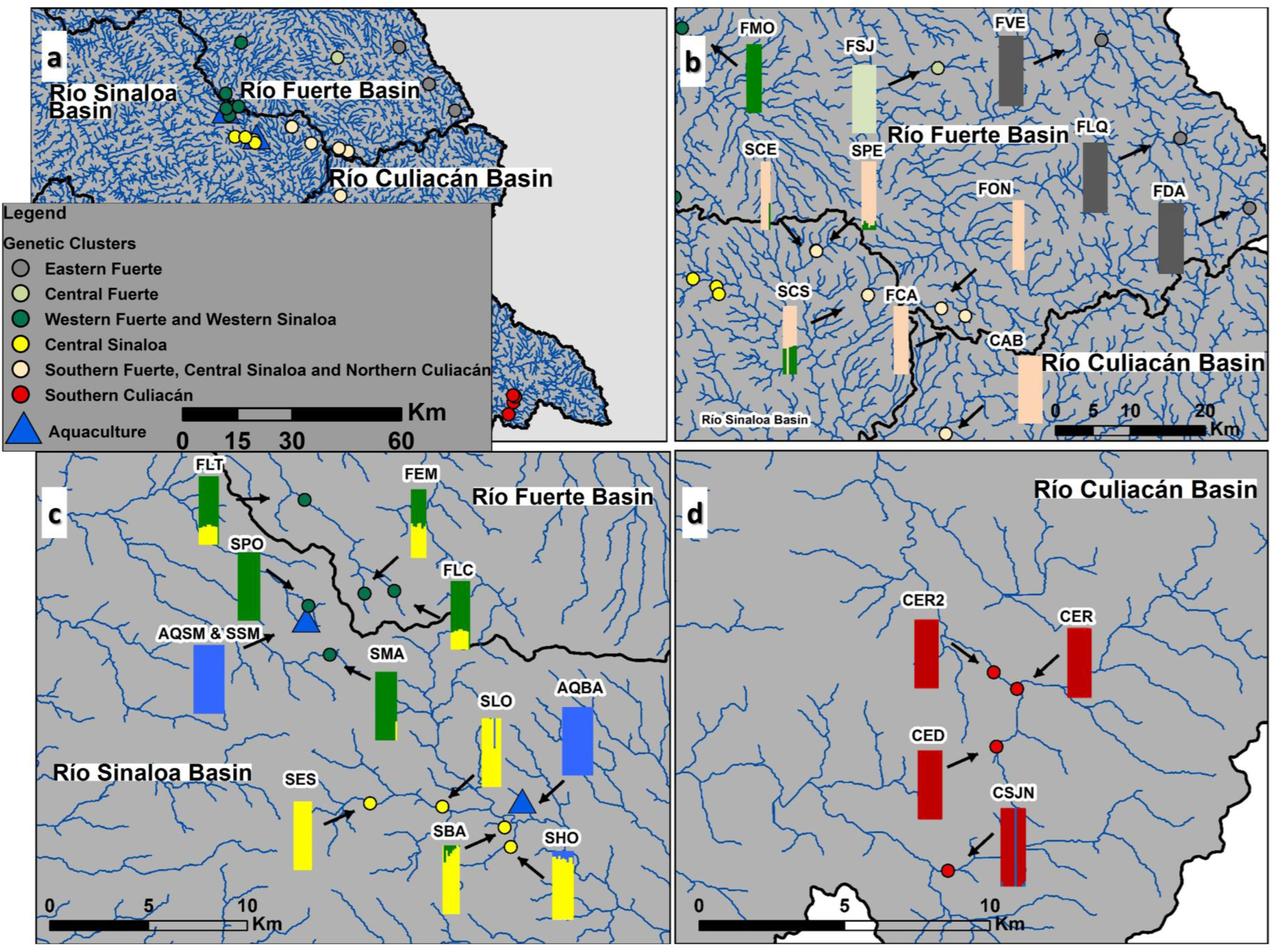
Spatial distribution of native *O. chrysogaster* and aquaculture rainbow trout genetic clusters defined in FastStructure. a) Spatial genetic structure across all the study area. b) Trout from Eastern Fuerte; Central Fuerte; Southern Fuerte, Central Sinaloa and Northern Culiacán genetic clusters; and FMO sample site from Western Fuerte and Western Sinaloa genetic cluster. c) Trout from Western Fuerte and Western Sinaloa; Central Sinaloa; and Aquaculture genetic clusters. d) Trout from Southern Culiacán genetic cluster. See table 1 for the explanation of the sample site codes.

### 3.3. Riverscape genetics analyses

Both latitude and precipitation of the driest month showed significant correlations with *H_E_*. These correlations reflect the negative effect of latitude and precipitation of the driest month on genetic diversity (Table 2).

**TABLE 2.**
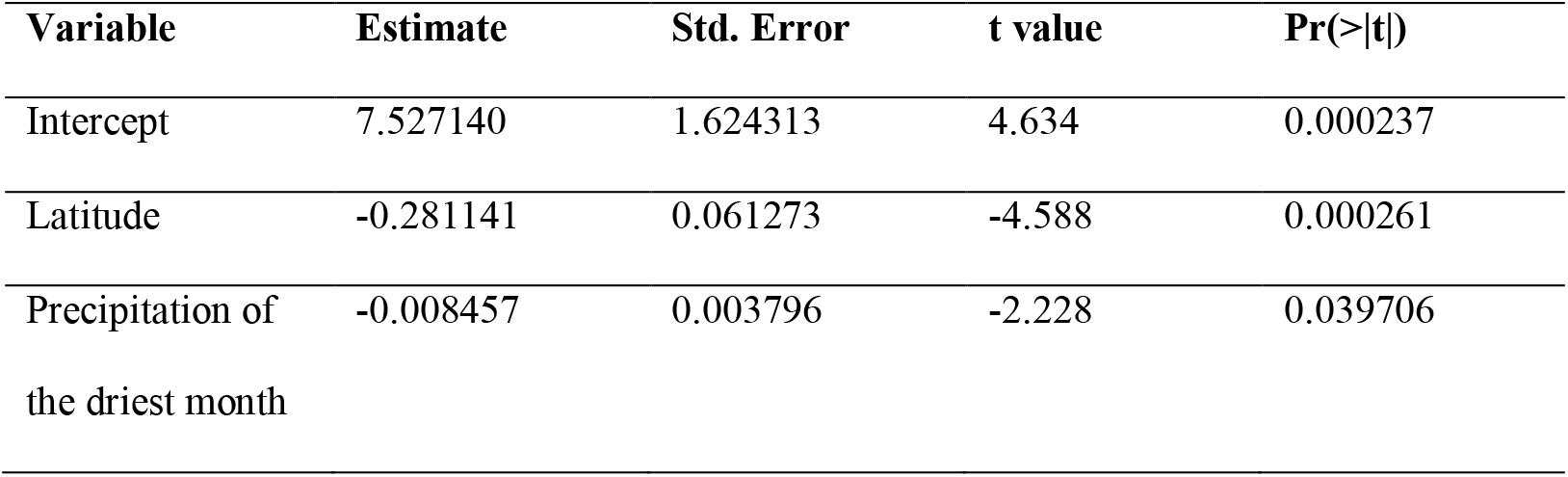
Generalized linear model explaining the effect of riverscape variables on expected heterozygosity (*H_E_*) for Mexican golden trout populations (Dataset C).

| Variable | Estimate | Std. Error | t value | Pr(>\|t\|) |
| --- | --- | --- | --- | --- |
| Intercept | 7.527140 | 1.624313 | 4.634 | 0.000237 |
| Latitude | -0.281141 | 0.061273 | -4.588 | 0.000261 |
| Precipitation of<br>the driest month | -0.008457 | 0.003796 | -2.228 | 0.039706 |

The partial Mantel test between pairwise *F_ST_*/(1 - *F_ST_*) and riverscape resistance at Río Sinaloa was highly statistically significant (p = 0.0002 and r = 0.87), suggesting that riverscape resistance strongly influences genetic divergence within this basin. Remarkably, the effect of riverscape resistance was not observed at Río Fuerte, probably due to the sampling strategy (Table 3). Is worth mentioning that at Río Sinaloa the sampling site distribution was restricted to a small area (few streams) along the length of the river, which is not the case for Río Fuerte.

**TABLE 3.**
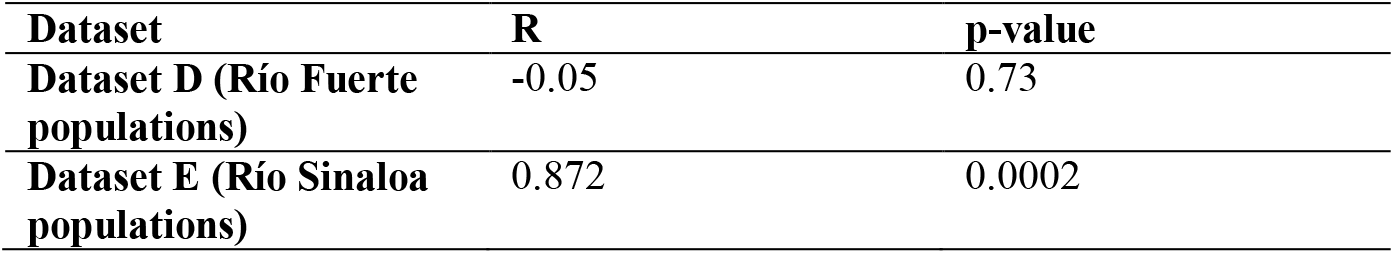
Partial Mantel test performed under 9,000 permutations between (*F_ST_*/(1-*F_ST_*) and riverscape resistance matrices.

### 3.4. Detection of SNPs under divergent selection

A total of 566 outlier loci were detected with PCAdapt, Bayenv2, and LFMM (Fig. 5). PCAdapt identified 278 outliers. On the other hand, Bayenv2 and LFMM identified 306 outliers correlated with environmental variables (temperature of the warmest quarter and precipitation of the driest month). Few outliers (96) were detected twice or more by the different approaches. No common outlier was shared among methods.

**Fig. 5.**
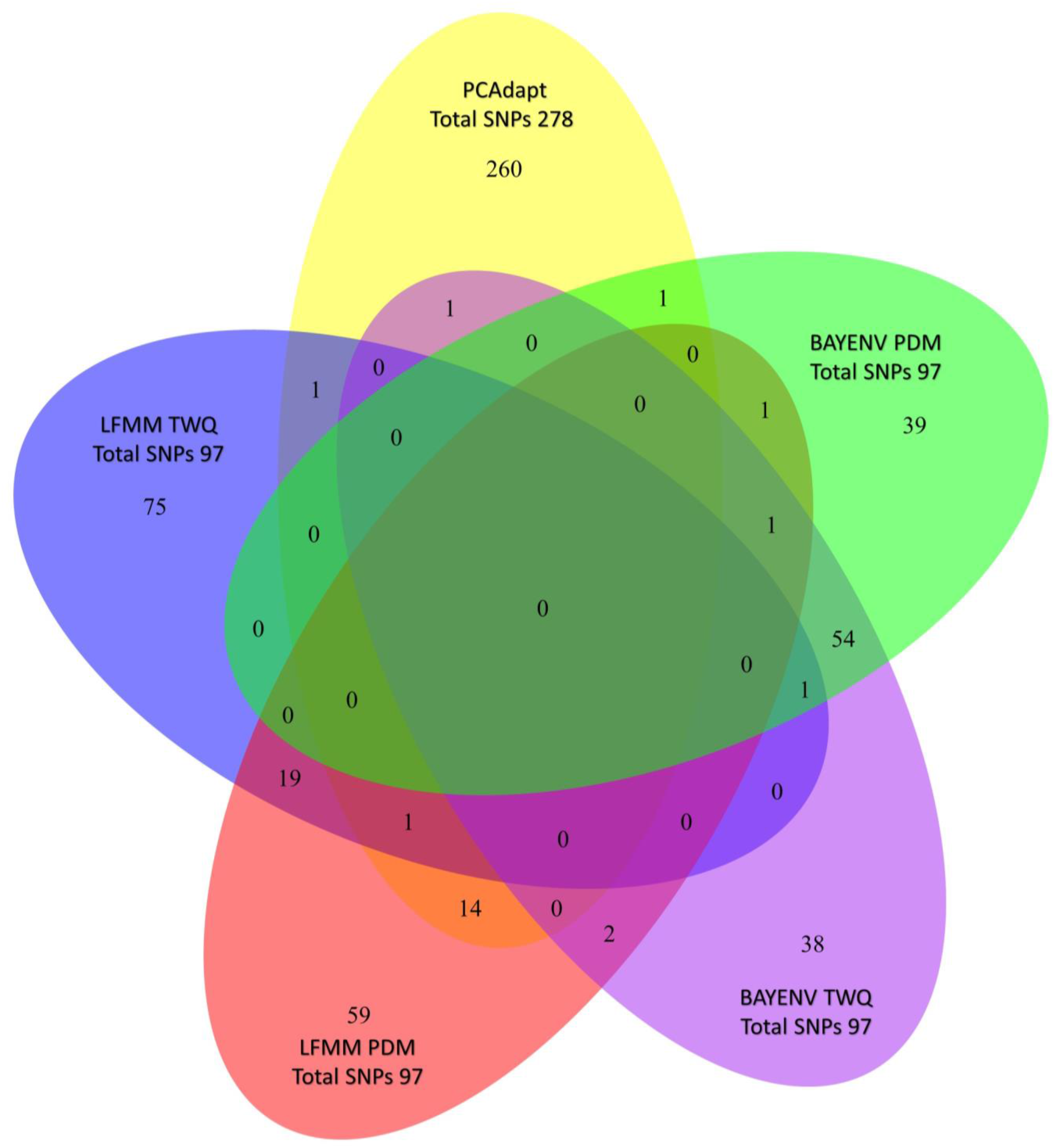
Venn diagram of outliers detected for *O. chrysogaster* among PCAdapt, LFMM using temperature of the warmest quarter (LFMM TWQ), LFMM using precipitation of the driest month (LFMM PDM), Bayenv2 using temperature of the warmest quarter (BAYENV TWQ), and Bayenv2 using precipitation of the driest month (BAYENV TWQ). The total amount of outliers (SNPs) detected by each method are represented in bold. Values outside of the curve intersections represent the number of outliers detected by only one method and the values inside the intersections represent the number of outliers detected by two methods or more.

After applying quality filters in the gene ontology analysis, 21 SNP loci under divergent selection and with protein annotations, were retained (Table 4). Most of these annotations were associated to biological functions in the literature (e.g. growth, reproduction and thermal tolerance; Table 4).

**TABLE 4.** Blast hits from sequences containing an SNP found to be putatively under selection by PCAdapt, BAYENV and LFMM. SNPs are identified by locus ID. Sequence names are identified as in Berthelot *et al.* (2014). Statistical significance of the hits is represented by the e-Value. Functional characterizations by gene ontology terms obtained in Blast2GO are identified by GO IDs.

| Locus ID | Sequence Name | e-Value | GO ID's | Known functions | Associated functions |
| --- | --- | --- | --- | --- | --- |
| CLocus_63824 | GSONMP00076296001 | 1.93E-08 | F:GO:0003676 | F:nucleic acid binding | Sex expression, lipid metabolism (Taggart et al. 2008; Hale et al. 2011) |
| CLocus_17128 | GSONMP00061816001 | 0.0000145 | F:GO:0008270;<br>P:GO:0030163 | F:zinc ion binding; P:protein catabolic process | Adaptation to elevated concentrations of waterborne (Hogstrand et al. 1994) |
| CLocus_24893 | GSONMP00073281001 | 0.000209 | F:GO:0003676 | F:nucleic acid binding | Sex expression, lipid metabolism (Taggart et al. 2008; Hale et al. 2011) |
| CLocus_34692 | GSONMP00068395001 | 0.00000574 | F:GO:0005524;<br>F:GO:0004672;<br>P:GO:0006468 | F:ATP binding; F:protein kinase activity; P:protein phosphorylation | Water flow and temperature acclimation (Milano et al. 2014) |
| CLocus_39212 | GSONMP00076439001 | 0.00000039 | P:GO:0097264;<br>C:GO:0005887;<br>F:GO:0005515;<br>P:GO:0007165;<br>C:GO:0016021;<br>F:GO:0005102 | C:integral component of plasma membrane; P:self proteolysis; P:signal transduction; F:receptor binding | Temperature adaptation (Crockett 1998) |
| CLocus_40408 | GSONMP00028710001 | 0.000684 | P:GO:0055085;<br>C:GO:0016021 | P:transmembrane transport; C:integral component of membrane | Temperature adaptation (Crockett 1998) |
| CLocus_40780 | GSONMP00020503001 | 0.000225 | F:GO:0005515 | F:protein binding | Water flow and temperature acclimation (Milano et al. 2014) |
| CLocus_41412 | GSONMP00076296001 | 0.000311 | F:GO:0003676 | F:nucleic acid binding | Sex expression, lipid metabolism (Taggart et al. 2008; Hale et al. 2011) |
| CLocus_45731 | GSONMP00008827001 | 0.000766 | F:GO:0008641 | F:small protein activating enzyme activity |  |
| CLocus_48136 | GSONMP00021876001 | 0.000665 | C:GO:0005615;<br>P:GO:0010506 | C:extracellular space; P:regulation of autophagy |  |
| CLocus_5089 | GSONMP00082580001 | 0.000619 | F:GO:0005515 | F:protein binding | Water flow and temperature acclimation (Milano et al. 2014) |
| CLocus_54656 | GSONMP00051628001 | 0.00000314 | F:GO:0005515 | F:protein binding | Water flow and temperature acclimation (Milano et al. 2014) |
| CLocus_55309 | GSONMP00044696001 | 0.000544 | F:GO:0003677;<br>C:GO:0005634;<br>F:GO:0008270; | C:nucleus; F:DNA binding; F:zinc ion binding; F:steroid hormone receptor activity; | Sex expression, lipid metabolism (Taggart et al. 2008; Hale et al. 2011) |
|  |  |  | F:GO:0003707;<br>P:GO:0006355;<br>P:GO:0043401 | P:regulation of transcription,<br>DNA-templated; P:steroid<br>hormone mediated signaling<br>pathway |  |
| CLocus_5644 | GSONMP00023521001 | 0.000247 | F:GO:0004725;<br>P:GO:0006470 | P:tyrosine metabolic process;<br>F:protein tyrosine phosphatase<br>activity; P:protein<br>dephosphorylation | Water flow and temperature acclimation<br>(Milano et al. 2014) |
| CLocus_58838 | GSONMP00030033001 | 0.00000662 | P:GO:0008360;<br>F:GO:0003779;<br>P:GO:0007010;<br>P:GO:0016043;<br>F:GO:0017048;<br>F:GO:0005488;<br>P:GO:0030036 | P:regulation of cell shape; F:actin<br>binding; F:Rho GTPase binding;<br>P:actin cytoskeleton organization | Water flow and temperature acclimation<br>(Milano et al. 2014) |
| CLocus_59529 | GSONMP00030319001 | 0.0000646 | P:GO:0016337;<br>F:GO:0005515;<br>P:GO:0007155;<br>F:GO:0005488;<br>P:GO:0032467;<br>P:GO:0043547 | P:single organismal cell-cell<br>adhesion; F:protein binding;<br>P:positive regulation of<br>cytokinesis; P:positive regulation<br>of GTPase activity | Water flow and temperature acclimation<br>(Milano et al. 2014) |
| CLocus_63824 | GSONMP00059101001 | 6.73E-07 | F:GO:0003676;<br>F:GO:0005524;<br>C:GO:0005634;<br>F:GO:0003677;<br>P:GO:0015074;<br>P:GO:0006355;<br>P:GO:0006313 | C:nucleus; F:DNA binding;<br>F:ATP binding; P:DNA<br>integration; P:regulation of<br>transcription, DNA-templated;<br>P:transposition, DNA-mediated | Sex expression, lipid metabolism (Taggart<br>et al. 2008; Hale et al. 2011) |
| CLocus_6876 | GSONMP00019151001 | 0.00052 | F:GO:0003676 | F:nucleic acid binding | Sex expression, lipid metabolism (Taggart<br>et al. 2008; Hale et al. 2011) |
| CLocus_69641 | GSONMP00059101001 | 0.0000108 | F:GO:0003676;<br>F:GO:0005524;<br>C:GO:0005634;<br>F:GO:0003677;<br>P:GO:0015074;<br>P:GO:0006355;<br>P:GO:0006313 | C:nucleus; F:DNA binding;<br>F:ATP binding; P:DNA<br>integration; P:regulation of<br>transcription, DNA-templated;<br>P:transposition, DNA-mediated | Sex expression, lipid metabolism (Taggart<br>et al. 2008; Hale et al. 2011) |
| CLocus_71994 | GSONMP00069497001 | 0.0000141 | F:GO:0005515;<br>F:GO:0005509 | F:protein binding; F:calcium ion<br>binding | Water flow and temperature acclimation<br>(Milano et al. 2014) |
| CLocus_77569 | GSONMP00045823001 | 0.0000424 | F:GO:0003676 | F:nucleic acid binding | Sex expression, lipid metabolism (Taggart et al. 2008; Hale et al. 2011) |

## 4. DISCUSSION

Our main objective was to explore the effect of both riverscape and cultured rainbow trout farm escapes, on the genetic diversity and local adaptation of an endemic salmonid. Low genetic admixture between aquaculture *O. mykiss* and native trout was found for the populations sampled. This study also revealed high genetic differentiation among geographically isolated locations, with populations in the south of the study area being the most isolated. A significant influence of latitude and precipitation of the driest month on genetic diversity was detected, in addition to evidence of isolation by riverscape resistance within a basin. Outlier detection and gene ontology analyses identified genes that could be implicated in adaptation to local climate heterogeneity. This integrative approach reveals that the riverscape influences different microevolutionary processes (i.e. gene flow and local adaptation) depending on the spatial scale: *i) at local scale:* genetic divergence might be shaped by hydrologic and topographic gradients and ruptures at intra-basin level; *ii) at regional scale:* temperature and precipitation influence genetic diversity within basins, but also among non-distant places at different basins; iii) at *broad scale:* temperature and precipitation gradients could be implicated in local adaptation. These findings, in addition to shedding light on riverscape genomics, are discussed in the context of the development of management strategies for endangered riverine species in order to preserve both genetic diversity and adaptive potential in the face of global change.

### 4.1. Absence of high levels of admixture between *O. chrysogaster* and aquaculture *O. mykiss*

The low genetic admixture (introgression) between native and exotic trout found in this study can be explained by the lack of proximity to aquaculture sites, as intensive aquaculture activities were not detected in the study area (authors’ field observations). Other studies at broader spatial scales found evidence of high genetic admixture, but mostly for undescribed Mexican trout forms in southernmost SMO areas, characterised by intensive aquaculture activities (Abadía-Cardoso et al. 2015; Escalante et al. 2014). In addition to reproductive behaviour, interspecific phenotypic variation and the overall geographic remoteness of aquaculture activities (Binder et al. 2015; Buchinger et al. 2017; Johnson et al. 2018); the low levels of exotic introgression found in this study might be explained by the local riverscape (i.e. altitude decrease; slope, stream order changes and temperature increase) acting as a boundary against exotic introgression. This physical barrier effect caused by the riverscape was suggested in recent simulation studies for this same species (Escalante et al. 2018), but also by empirical data for *O. mykiss*, *O. clarkii lewisi*, *O. clarkii bouvieri* and *Salmo trutta* (Weigel et al. 2003; Gunnell et al. 2008; Splendiani et al. 2013). The extensive introduction of exotic trout has been documented as one of the greatest threats to the entire endemic Pacific trout complex (Miller et al. 1989; Bahls 1992; Penaluna et al. 2016). Also, the harmful effects of exotic salmonid invasions have been recognised in native salmonid populations all over the world through parasites, native food web alterations, competition, and replacement of the native gene pool (Heggberget et al. 1993; Fausch 2007; Muhlfeld et al. 2009; Marie et al. 2012; Vera et al. 2017). In the SMO, particularly, due to economic interests, it is expected that aquaculture activities will increase in coming years, while their impact on native trout populations is still little known, making it challenging to implement management plans for conservation purposes (Hendrickson et al. 2002).

However, the low level of admixture between native and exotic trout detected in this study highlights the beneficial effect of environmental heterogeneity and low aquaculture activity, but would inevitably be reversed if this activity increases. Therefore, the proper use of native trout for aquaculture activities in the area only using closely-related populations and respecting the ecological niches, may offer an alternative which avoids the harmful effects of exotic trout invasions. The low exotic admixture with aquaculture trout observed in *O. chrysogaster* in previous works supports our results. Indeed, in the specific case of Arroyo Agua Blanca in Río Culiacán, previous studies reported high levels of genetic admixture with *O. mykiss* in an analysis of wild samples collected in 1997 (Escalante et al. 2014), while samples collected in 2015 used in this study did not display genetic admixture. These results are encouraging and revealed that the genetic pool of riverine species could still be preserved despite the existence of aquaculture practices in their distribution area. Thus, conservation strategies may be considered to avoid the probable harmful effects of exotic introgression in *O. chrysogaster* populations, such as banning or strongly regulating rainbow trout aquaculture.

### 4.2. Riverscape drivers of genetic diversity

Low levels of heterozygosity were correlated with an increase in latitude, and precipitation of the driest month. The genetic diversity values observed here are lower than those reported for trout from northern latitudes (e.g. Carim et al. 2016; Linløkken et al. 2016; Wenne et al. 2016; Perreault-Payette et al. 2017; Winans et al. 2018; Pearse and Campbell 2018), this is in agreement with former studies that already showed the lower genetic diversity of *O. chrysogaster* in relation to trout form USA (Escalante et al. 2014; Abadía-Cardoso et al. 2015). From an ecological point of view, the main sources of variation in genetic diversity are*: i) variation in effective size*: populations with large effective sizes are expected to have higher heterozygosity than populations with smaller effective sizes, as they have a larger number of breeders that keep allele frequencies stable (Höglund 2009); *ii) space, mainly through variation in the environment and habitats:* isolated populations with low immigration rates may exhibit inbreeding and reduced gene flow with closely related populations, as they are exposed to genetic drift (Riginos and Liggins 2013); iii) *contemporary and historical events, and human disturbances*: the fragmentation or connection of populations with consequent variations in genetic diversity due to microevolutionary processes (e.g. gene flow and local adaptation) (Hewitt 2000; Banks et al. 2013).

The occurrence of extreme hydroclimatic events during embryo incubation may have catastrophic consequences on trout populations, causing strong variations in population sizes (Hand et al. 2016), as well as variations in the effect of genetic drift. In this study, a negative correlation was detected between precipitation of the driest month on *H_E_*. This correlation may be explained by the fact that *O. chrysogaster* mates during the dry season (December – March; García-De León et al. 2016). Flash flood events during the reproductive period of *O. chrysogaster* may cause embryo mortality and, consequently, dramatic reductions in population size that translate into a decline in heterozygosity. A negative influence of precipitation on heterozygosity has also been observed in native steelhead trout in the USA, with similar life history traits and habitat conditions to *O. chrysogaster* (Narum et al. 2008).

A negative influence of latitude in *H_E_* was also observed. The heterozygosity is actually lower on the periphery of the study area (i.e. east of Río Fuerte, southwest of Río Fuerte, northwest of Río Sinaloa and south of Río Culiacán) (Fig. 2). These peripheral zones have been defined in previous studies at the limits of the species’ ecological niche (Ruiz-Luna et al. 2017; Escalante et al. 2018). This pattern of genetic diversity deficit on the periphery due to isolation has been broadly observed in a wide number of species (de Lafontaine et al. 2018). Taken together, the low genetic diversity, high *F_ST_* values, geographical remoteness, high riverscape resistance (observed in riverscape resistance surfaces) and habitat suitability observed in previous studies using ecological niche models (Escalante et al. 2018) suggest that populations in the south of Río Culiacán (i.e. CED, CER, CER2 and CSJN) are the most isolated in this study.

The small *N_E_* found in our study (≤146) fall far short of what is needed (≥ 500) to preserve the long-term viability of salmonids, which raises the question of the long-term survival of the populations studied (Koskinen et al. 2002; Rieman and Allendorf, 2001). Moreover, the low values of *N_E_* (≤ 500) could be the reason for the lack of a positive correlation between the *N_E_* and *H_E_* reported in this study. Certainly, the effects of drift are shown in sites with high *N_E_* values and low *H_E_* values and vice versa, for example FLQ had *N_E_* of 116.8 and *H_E_* of 0.05, while FCA had *N_E_* of 8.2 and *H_E_* of 0.13 (Table 1). However, generally low *N_E_* and *H_E_* values are common in species that experience rapid expansions after glaciation periods because of habitat shifts and isolation (Hewitt 2000). This decline in genetic diversity may therefore be due to a combination of several factors: bottlenecks occurring after *O. chrysogaster* colonization of the SMO at the end of the Pleistocene (Behnke et al. 2002), and/or habitat fragmentation due to riverscape resistance impeding immigrant exchange among populations (Channell and Lomolino 2000; Behnke et al. 2002; Eckert et al. 2008; Ruzzante et al. 2016). Increasing the sample size in further studies would make it possible to test these hypotheses.

The highest parts of SMO inhabited by *O. chrysogaster* should be made into natural protected areas with strictly regulated land use and land change activities. The aforementioned strategy can help to preserve dispersal corridors among populations maintaining the gene flow but also habitat quality, which is necessary to ensure the native genetic diversity (Olsen et al. 2017). Moreover, under any past or present scenarios explaining the low levels of *N_E_*, avoiding the loss of genetic diversity should be considered for the development of conservation strategies. An alternative that could be explored in depth is the translocation of individuals between small and isolated populations containing the same gene pool, to preserve the genes involved in local adaptation (Sato and Harada, 2008). Translocation is a conservation strategy that can help to avoid genetic diversity reduction, and could be achieved by ensuring founding populations including as many individuals as possible, and augmenting the relocated populations after establishment with new trout from existing populations (Faulks et al. 2017). However, this strategy requires extensive genetic monitoring of both existing and relocated populations in order to avoid outbreeding problems.

### 4.3. Riverscape drivers of genetic divergence

The influence of riverscape structure on genetic divergence has already been tested in salmonids, with riverscape defined as a fundamental driver of gene flow (Kanno et al. 2011; Torterotot et al. 2014; Landguth et al. 2016). In this study, partial Mantel tests suggest an influence of riverscape resistance in genetic variation only in Río Sinaloa. These outputs might be due to the sampling strategy, which was closer to a least cost path approach in Río Sinaloa with the sampling sites homogeneously distributed along a few streams on a small spatial scale, compared with Río Fuerte where the sampling sites are distributed over a larger spatial extent in several streams. Demo-genetic simulations have already suggested that riverscape resistance has a strong influence on the genetic divergence of *O. chrysogaster* at Río Fuerte (Escalante et al. 2018). Therefore, we expect that the effect of the riverscape at Río Fuerte can be evidenced with a more continuous sampling strategy along the length of the streams (Cushman and Landguth 2010). Moreover, the use of a point based approach to test the effect of local riverscape conditions on genetic divergence might be considered to better understand the influence of environmental factors on the species’ genetic structure (Grummer et al. 2019).

The effect of riverscape resistance on gene flow may determine adaptation to local environments, due to the fact that gene flow allows the entry of new genes, and then, through recombination during sexual reproduction, a new combination of genes is produced and populations gain genetic advantages to cope with changing environments (Kokko et al. 2017). For endangered species living in restricted heterogeneous habitats, it is vital to understand the relationship between riverscape and gene flow in order to define management units based on populations with historical gene flow, and which occupy similar ecological niches (Crandall et al. 2000; Olsen et al. 2017; Schmidt et al. 2017). It is important to define the influence of landscape factors on gene flow processes among populations at local scales (within basins), to then understand adaptive variation across wider scales (along different basins). Thus, the development of riverscape resistance surfaces could help the conservation of native trout, defining and preserving dispersal corridors to maintain gene flow among closely related populations with low genetic diversity.

### 4.4. Detection of SNPs under divergent selection

The 306 SNPs correlated with hydroclimatic variables (i.e. precipitation of the driest month and temperature of the warmest quarter), suggest that these environmental variables may act as selective factors in *O. chrysogaster*. The influence of temperature and precipitation on adaptive genetic variation has also been suggested for steelhead trout from the Inner Columbia River Basin and for cutthroat trout from the Great Basin Desert both in U.S.A. (Hand et al. 2016; Amish et al. 2019). Temperature has been defined as a potential driver of adaptive processes in salmonids mainly because ectothermic body temperature is closely associated with the environment (Hecht et al. 2015; Hand et al. 2016). Moreover, variations in temperature and precipitation affect phenomena such as changes in dispersal and reproduction timing, age at maturity, growth, fecundity, and survival (Crozier & Hutchings, 2014; Hecht et al., 2015). The very low overlap of outliers observed by the different approaches used in this study is expected when using different methods, since their algorithms and prior assumptions differ considerably (Ahrens et al. 2018; Dalongeville et al. 2018; Amish et al. 2019).

Among the annotated gene functions detected by a gene ontology analysis in outlier loci, actin-binding proteins might play an important role in water flow and temperature acclimation, as reported for other species such as the European hake (Milano et al. 2014). Thus, adaptive genetic variation across the study area may increase among populations exposed to different temperature and precipitation regimes. However, analyses of adaptive variation, using the population genetics analyses of outlier loci, are needed to confirm this. Moreover, protein binding is also associated with trout growth and flesh quality, which in turn may be associated with stream flow and water temperature (Salem et al. 2010). Even though our results suggest potential adaptation of *O. chrysogaster* to local conditions, these findings should be interpreted with caution. It is acknowledged that when sample size is small and the averaged genetic differentiation is high, this may result in false positive gene-environment associations (Hoban et al. 2016).

Our findings, together with previous works on related species, indicate that salmonids develop adaptations to cope with changing climatic conditions (Hecht et al. 2015; Bourret et al. 2013). However, faced with the threat of man-made global warming, habitat fragmentation and exotic introduction the species’ risk of extinction is high, especially for small populations with putative fitness loss. Indeed, actions at local scale may not be enough to ensure the survival of the species (Rahel et al. 2008; Lawler et al. 2010). On this basis, federal agencies should consider climate change in all their management plans, integrating multidisciplinary approaches to understand the effect of climate fluctuations on native species at different scales. The landscape genetics approach considered in this study may be helpful in the development of these management strategies and illustrates how genomic data can inform monitoring programs and conservation actions (Flanagan et al. 2018; Hendricks et al. 2018). Finally, it is a matter of urgency that predictive models associating climate change with genomic diversity in order to maintain local adaptation skills are used to design the best conservation strategies for all endangered salmonid species in North America. This broad evolutionary perspective is crucial for the application of adaptive genomic variation to conservation efforts (Pearse 2016; Leitwein et al. 2017; Razgour et al. 2019). However, as in many other Latin American countries, until now Mexican federal agencies showed a lack of interest to invest financial resources and integrate genomic tools into conservation strategies (Torres-Florez et al. 2017), making more challenging the preservation of native salmonids in relation to USA and Canada. Therefore, Government authorities should understand the importance of the incorporation of evolutionary research and management actions, linking theory and practice in order to ensure the native biodiversity in the face of global change. Undoubtedly, improving this requires improving connectivity between scientists and policy makers (Torres-Florez et al. 2017).

### 4.5. Conclusions

The integrative study presented here describes the effect of both landscape and anthropogenic factors on microevolutionary processes providing information about possible effects of hydroclimatic variables on local adaptation. This type of approach has not been widely considered in conservation genomics studies, especially in Latin America. Our findings suggest that both low genetic introgression, and that riverscape factors have an effect on genetic diversity, connectivity and adaptive genetic variation. The approach presented here may be useful for developing conservation strategies for endangered riverine fish species that consider both the ecological and evolutionary aspects.

## ACKNOWLEDGEMENTS

This research was supported under projects funded by The Mexican Council of Science and Technology (CONACYT; Ref. CB-2010-01-152893) and ECOS Nord (M14A04). Marco A. Escalante was a recipient of a Mexican Council of Science and Technology fellowship for his PhD studies. We are especially grateful to R. Hernández G., L. A. Ramirez H., and other members of CIAD staff for all their support in the planning, logistics and conducting of field surveys. We are grateful to G. Ingle from the Mexican Fisheries Institute (INAPESCA) for facilitating the obtention of trout samples. Thanks also to T. Durán for his support during field surveys. We would like to thank all the local guides who helped us during trout sampling, especially Ricardo Silvas ‘’El Chalote’’. We are also grateful to the municipal government of Guanaceví, PRONATURA, and WWF for their assistance with field surveys. Thanks to all members of the ‘’Truchas Mexicanas’’ binational group, and especially G. Ruiz C. and F. Camarena R. Thanks to B. Shepard, C. Kruse, J. Dunham, and others for donating electrofishing equipment. We also wish to thank F. Valenzuela Q., C. A. Rivera, A. Delongeville and the staffs of the Molecular Markers Platform at the CEFE and the Laboratorio de Genética para la Conservación at CIBNOR for their support with the molecular biology work. Sampling was conducted under the collection permit numbers SGPA/DGVS/02485/13 and SGPA/DGVS/05052/15.

## SUPPORTING INFORMATION

Additional Supporting Information can be found in the online version of this article:

**Online Resource 1.** Six datasets on four different spatial scales.

**Online Resource 2.** Riverscape resistance surfaces and matrices.

**Online Resource 3.** *F_ST_* coefficients among sample sites of Mexican golden trout, aquaculture rainbow trout and lab hybrids.

**Online Resource 4.** Assignment probabilities obtained in fastStructure for different K values.

**Online Resource 5.** Ancestry coefficients obtained in faststructure for Dataset B with native trout without significant introgression.

## Online Resource 1

Five datasets on four different spatial scales.

Five datasets at four different spatial scales were constituted after quality filters (Table S1.1). Dataset A: to perform population genetics analyses, all samples after bioinformatics filtering were kept (native, aquaculture and hybrids) including 272 genotyped individuals (Table S1.2). Dataset B: To conduct landscape genomics analyses (gene-environment associations) individuals with aquaculture ancestry coefficients ≥ 20% (based on fastStructure outputs) were removed, the trout collected at Arroyo San Miguel (SSM) was discarded since it is close to aquaculture facilities. Aquaculture rainbow trout was also discarded. Similarly, one individual from Arroyo La Osera (SLO) and one individual from Arroyo San Juan del Negro (CSJN) who presented significant aquaculture ancestry coefficients in fastStructure outputs were not considered either (Table S1.3). Dataset C: In order to conduct landscape genetics analyses with spatially continuous populations, trout from the southernmost part of Río Culiacán was discarded from Dataset B to constitute this new dataset (Table S1.4). Additionally, riverscape genomics analyses were performed at basin scale using two datasets. Dataset D: including trout from 7 sampling sites at Río Fuerte Basin, three sampling sites (FDA, FLQ and FFVE) at the easternmost part of Río Fuerte were discarded due to their large riverine isolation (long riverine distance) in relation with the other sampling sites (Table S1.5). Dataset E: including non-significantly introgressed native trout from the 9 sampling sites at Río Sinaloa Basin (Table S1.6). Native trout from Río Culiacán Basin were not analyzed independently at basin scale due to the small amount of sample sites.

**Table S1.1.**
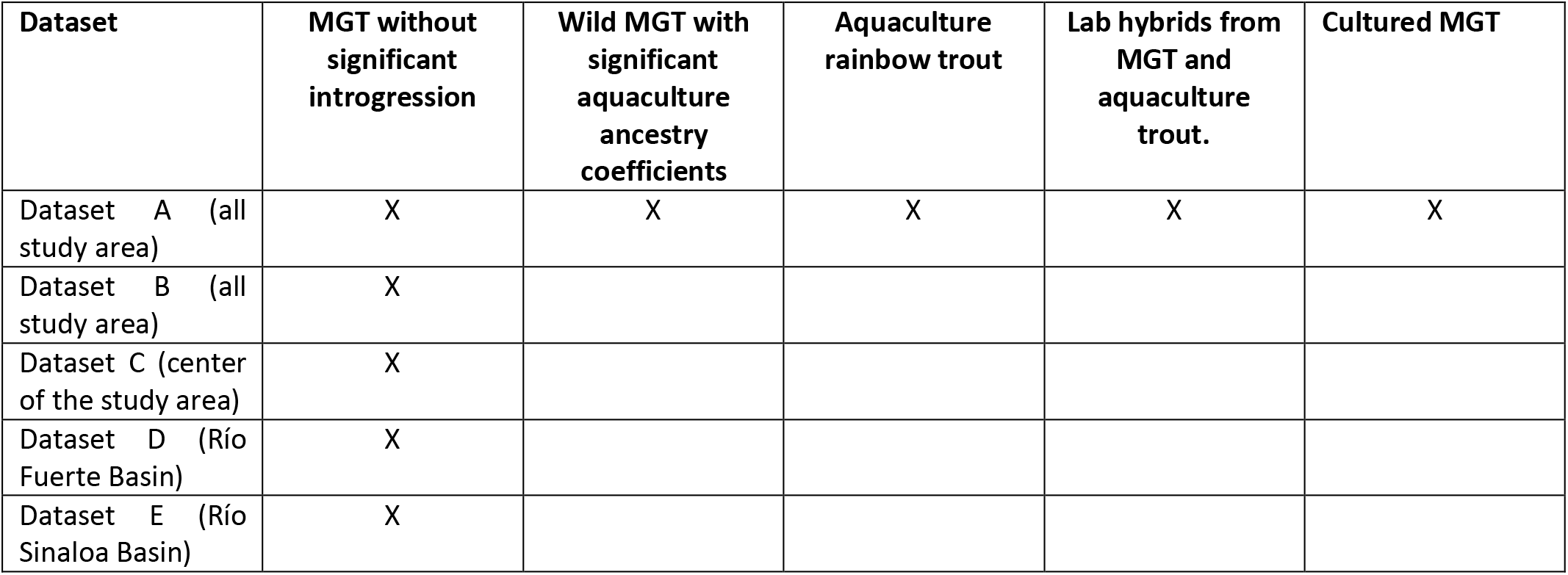
Characteristics of the six genetic datasets. (MGT. Mexican golden trout)

**Table S1.2.** Dataset A.

| Río Fuerte |  |  |  |  |  |
| --- | --- | --- | --- | --- | --- |
| CODE | Latitude | Longitude | Locality | Description | N |
| FDA | 26.07 | -106.31 | Arroyo del Agua | Wild trout | 10 |
| FEM | 26.09 | -107.02 | Arroyo El Manzano | Wild trout | 8 |
| FLC | 26.09 | -107.00 | Arroyo Las Cuevas | Wild trout | 10 |
| FLQ | 26.16 | -106.40 | Arroyo La Quebrada | Wild trout | 12 |
| FLT | 26.13 | -107.04 | Arroyo Las Truchas | Wild trout | 8 |
| FMO | 26.29 | -106.99 | Arroyo Momorita | Wild trout | 5 |
| FSJ | 26.24 | -106.69 | Arroyo San José | Wild trout | 12 |
| FCA | 25.94 | -106.65 | Arroyo Calera | Wild trout | 9 |
| FON | 25.95 | -106.68 | Arroyo La Onza | Wild trout | 6 |
| FVE | 26.28 | -106.49 | Río Verde | Wild trout | 12 |
| **Total** | - | - | - | - | **92** |
| Río Sinaloa |  |  |  |  |  |
| CODE | Latitude | Longitude | Locality | Description | N |
| SBA | 25.98 | -106.95 | Arroyo Baluarte | Wild trout | 9 |
| SES | 25.99 | -107.01 | Arroyo El Soldado | Wild trout | 9 |
| SHO | 25.97 | -106.95 | Arroyo Hondo | Wild trout | 11 |
| SLO | 25.99 | -106.98 | Arroyo La Osera | Wild trout | 10 |
| SMA | 26.06 | -107.03 | Arroyo Macheras | Wild trout | 10 |
| SPO | 26.08 | -107.04 | Arroyo Potrero | Wild trout | 11 |
| SSM | 26.07 | -107.04 | Arroyo San Miguel | Cultured trout | 4 |
| SCS | 25.97 | -106.77 | Arroyo Cerro Solo | Wild trout | 9 |
| SCE | 26.02 | -106.83 | Arroyo Cebollín | Wild trout | 7 |
| SPE | 26.02 | -106.83 | Arroyo Pericos | Wild trout | 7 |
| **Total** | - | - | - | - | **87** |
| Río Culiacán |  |  |  |  |  |
| CODE | Latitude | Longitude | Locality | Description | N |
| CED | 25.14 | -106.13 | Arroyo El Desecho | Wild trout | 12 |
| CER | 25.16 | -106.12 | Arroyo El Río 1 | Wild trout | 12 |
| CER2 | 25.17 | -106.13 | Arroyo El Río 2 | Wild trout | 12 |
| CSJN | 25.10 | -106.14 | Arroyo San Juan del Negro | Wild trout | 12 |
| CAB | 25.80 | -106.68 | Arroyo Agua Blanca | Wild trout | 10 |
| **Total** | - | - | - | - | **58** |
| Lab Trout |  |  |  |  |  |
| CODE | Latitude | Longitude | Locality | Description | N |
| H |  |  | INAPESCA Aquaculture Center | Hybrids from MGT and rainbow trout | 4 |
| REP |  |  | INAPESCA Aquaculture Center | Cultured MGT | 7 |
| **Total** | - | - | - | - | **11** |
| **Aquaculture farms** |  |  |  |  |  |
| **CODE** | **Latitude** | **Longitude** | **Locality (farm)** | **Description** | **N** |
| AQEB | 25.99 | -106.95 | El Barro Aquaculture | rainbow trout | 14 |
| AQSM | 26.07 | -107.04 | San Miguel Aquaculture | rainbow trout | 10 |
| **Total** | - | - | - | - | **24** |
| **Dataset total** |  |  |  |  | **272** |

**Table S1.3.**
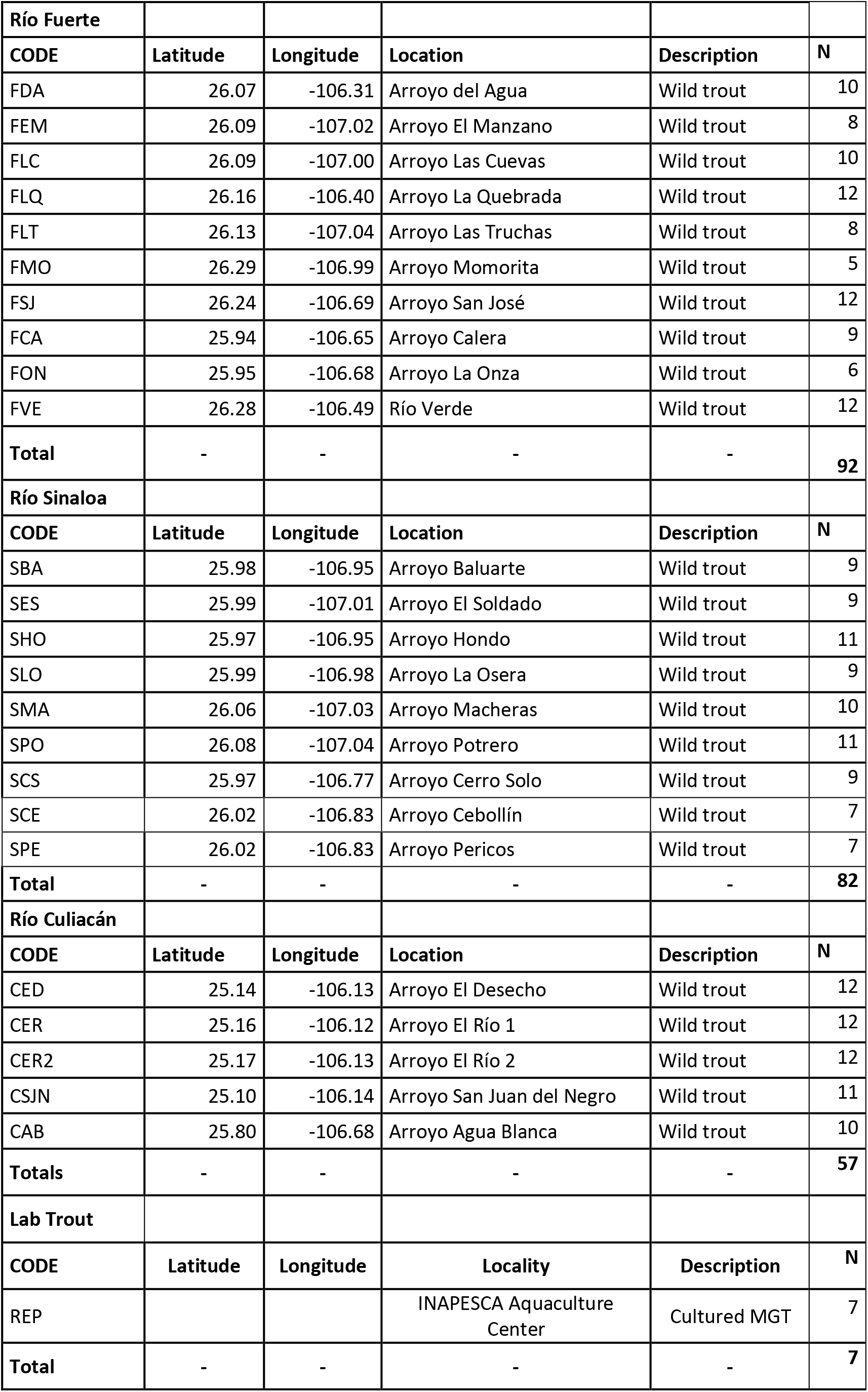
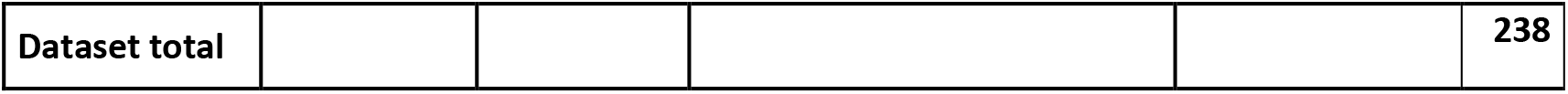
Dataset B.

**Table S1.4.**
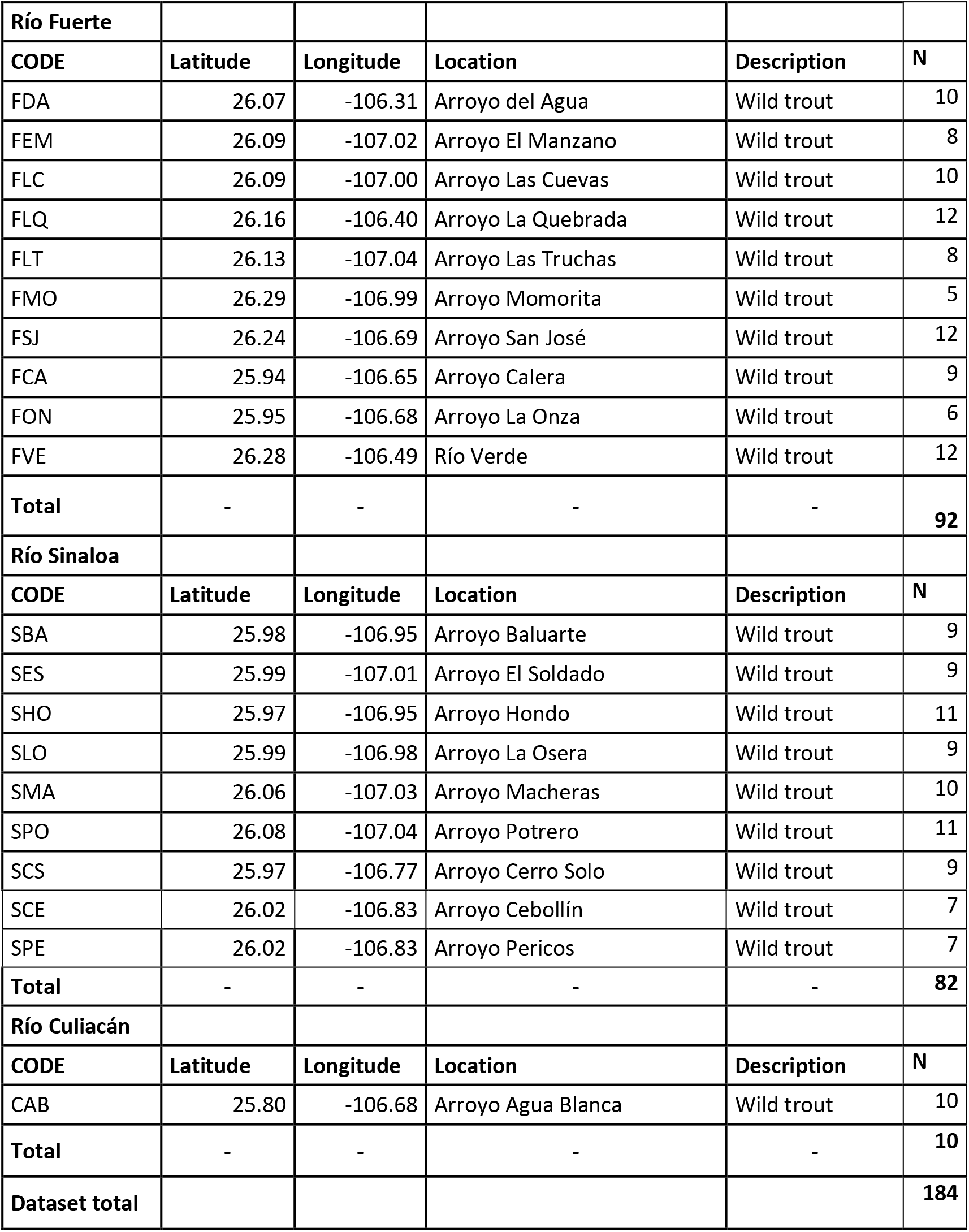
Dataset C.

**Table S1.5.** Dataset D.

|  |  |  |  |  |  |
| --- | --- | --- | --- | --- | --- |
| **Río Fuerte** |  |  |  |  |  |
| **CODE** | **Latitude** | **Longitude** | **Location** | **Description** | **N** |
| FLC | 26.09 | -107.00 | Arroyo Las Cuevas | Wild trout | 10 |
| FLT | 26.13 | -107.04 | Arroyo Las Truchas | Wild trout | 8 |
| FMO | 26.29 | -106.99 | Arroyo Momorita | Wild trout | 5 |
| FSJ | 26.24 | -106.69 | Arroyo San José | Wild trout | 12 |
| FCA | 25.94 | -106.65 | Arroyo Calera | Wild trout | 9 |
| FON | 25.95 | -106.68 | Arroyo La Onza | Wild trout | 6 |
| FVE | 26.28 | -106.49 | Río Verde | Wild trout | 12 |
| **Dataset total** | - | - | - | - | **62** |

**Table S1.6.** Dataset E.

| **Río Sinaloa** |  |  |  |  |  |
| --- | --- | --- | --- | --- | --- |
| **CODE** | **Latitude** | **Longitude** | **Location** | **Description** | **N** |
| SBA | 25.98 | -106.95 | Arroyo Baluarte | Wild trout | 9 |
| SES | 25.99 | -107.01 | Arroyo El Soldado | Wild trout | 9 |
| SLO | 25.99 | -106.98 | Arroyo La Osera | Wild trout | 11 |
| SHO | 25.97 | -106.95 | Arroyo Hondo | Wild trout | 9 |
| SMA | 26.06 | -107.03 | Arroyo Macheras | Wild trout | 10 |
| SPO | 26.08 | -107.04 | Arroyo Potrero | Wild trout | 11 |
| SCS | 25.97 | -106.77 | Arroyo Cerro Solo | Wild trout | 9 |
| SCE | 26.02 | -106.83 | Arroyo Cebollín | Wild trout | 7 |
| SPE | 26.02 | -106.83 | Arroyo Pericos | Wild trout | 7 |
| **Dataset total** | - | - | - | - | **82** |

## Online Resource 2

Riverscape resistance surfaces and matrices.

To analyze connectivity among Mexican golden trout populations, a riverscape resistance surface was created. Four environmental variables were selected (altitude, slope, temperature of the warmest quarter and stream order). Using a raster format (30 m pixel size) values from 0 to 10 were assigned to the pixels by variable based on the criteria reported by Escalante et al. (2018). Then, a resistance surface was produced averaging the values for the four variables (Table 1).

For subsequent analyses with the gdistance package (van Etten 2012), the output resistance values were rescaled between 1 and 2, the minimum or absence of riverscape resistance is represented by 1, while 2 is the maximum resistance. All those analyses were conducted in ArcGIS v 10.2 (ESRI 2013).

**Table S2.1.** Resistance values for riverscape variables: altitude, slope, temperature of the warmest quarter (TWQ) and stream order (SO).

| Altitude (m) | Slope (° Slope) | TWQ (°C) | SO | Resistance<br>value |
| --- | --- | --- | --- | --- |
| 3009 - 1950 | 0 - 6.74 | ≤ 14 | 1 - 3 | 0 |
| 1949 - 1900 | 6.75 - 13.48 | 15 |  | 1 |
| 1899 - 1850 | 13.49 - 20.22 | 16 |  | 2 |
| 1849 - 1800 | 20. 23 - 26.96 | 17 | 4 | 3 |
| 1799 - 1750 | 26.97 - 33.70 | 18 |  | 4 |
| 1749 - 1700 | 33.71 - 40.44 | 19 |  | 5 |
| 1699 - 1650 | 40.45 - 47.18 | 20 | 5 | 6 |
| 1649 - 1600 | 47.19 - 53.92 | 21 |  | 7 |
| 1599 - 1550 | 53.93 - 60.66 | 22 | 6 | 8 |
| 1549 - 1500 | 60.67 - 67.40 | 23 |  | 9 |
| ≤ 1499 | 67.41 - 74.14 | ≥ 24 | 7 | 10 |

To calculate isolation by riverscape resistance and linear distance, four matrices among sample sites within basins were generated in gdistance R package (van Etten, 2012): Isolation by riverscape resistance among Dataset D for Río Fuerte populations (Matrix I), Isolation by riverscape resistance among Dataset E for Río Sinaloa populations (Matrix II), Isolation by distance among Dataset D for Río Fuerte populations (Matrix III), and Isolation by distance among Dataset E for Río Sinaloa populations (Matrix IV). Matrix I and Matrix II were calculated using the riverscape resistance surface, while Matrix III and Matrix IV were calculated taking into account only the geographical coordinates of the sampling sites (Table S2.2).

**Table S2.2.**
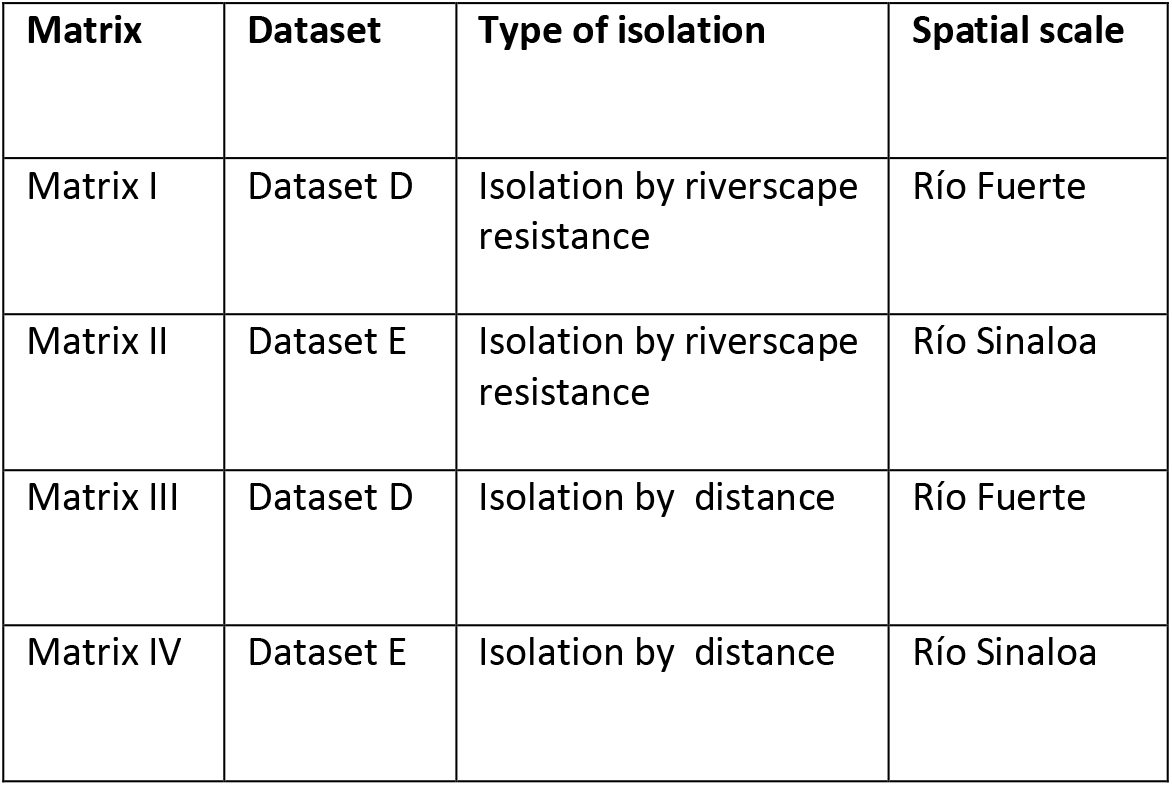
Isolation by distance and riverscape resistance matrices.

## Online Resource 3

*F_ST_* coefficients among sample sites of Mexican golden trout.

**Table S3.1.**
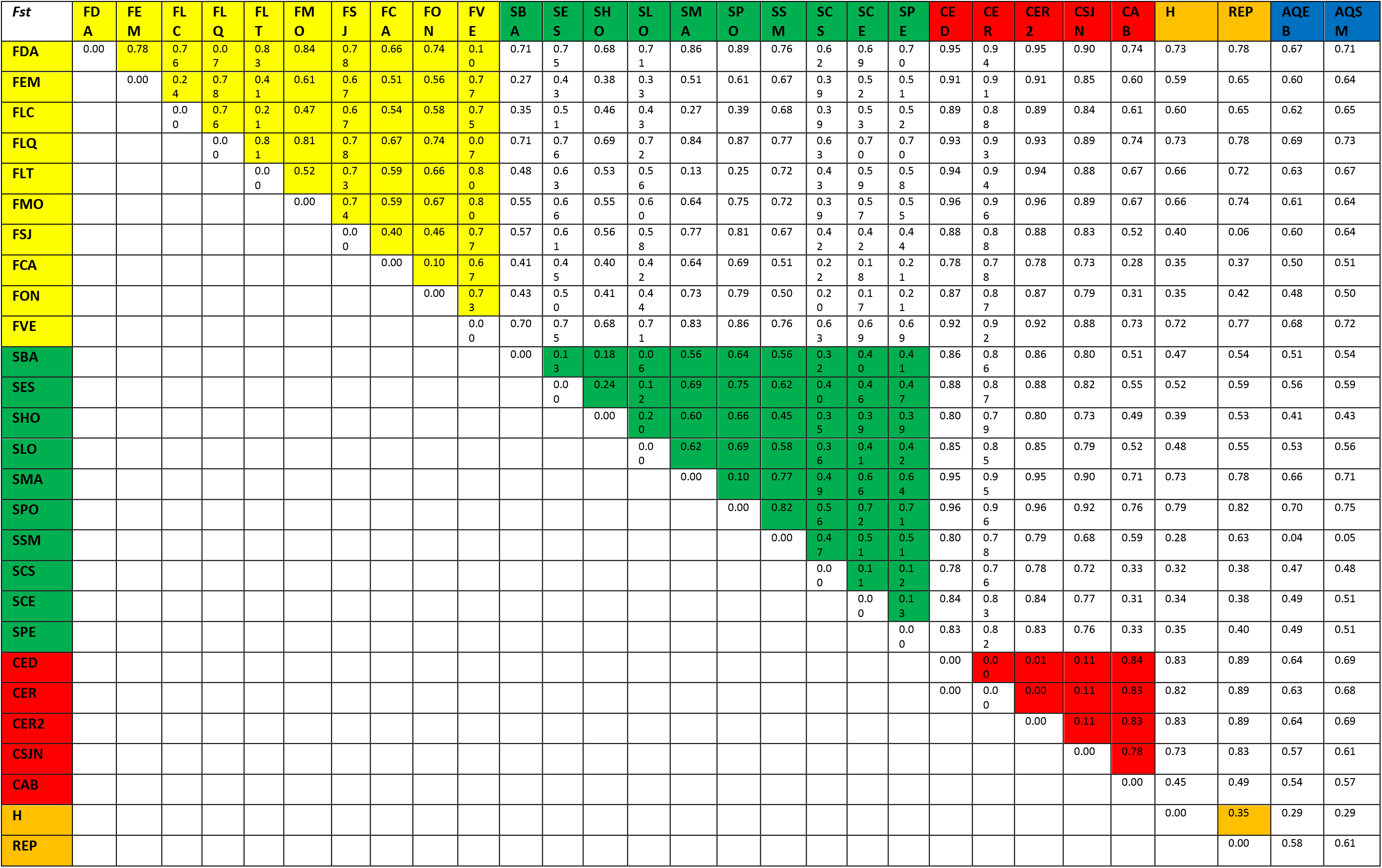
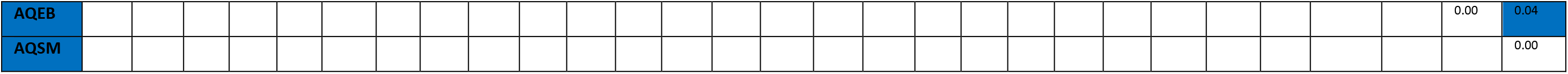
*F_ST_* coefficients among sample sites of Mexican golden trout, aquaculture rainbow trout and lab hybrids. Each sample site is represented by a code, including the initial of the river basin (F. Río Fuerte; S. Río Sinaloa; C. Río Culiacán) and an acronym of the locality Spanish name: Arroyo del Agua (FDA), Arroyo El Manzano (FEM), Arroyo Las Cuevas in Río Fuerte (FLC), Arroyo La Quebrada (FLQ), Arroyo Las Truchas (FLT), Arroyo Momorita (FMO), Arroyo San José (FSJ), Arroyo Caleras (FCA), Arroyo La Onza (FON), Arroyo Río Verde (FVE), Arroyo Baluarte (SBA), Arroyo El Salto (SES), Arroyo Hondo (SHO), Arroyo La Osera (SLO), Arroyo Macheras (SMA), Arroyo El Potrero (SPO), Arroyo San Miguel (SSM), Arroyo Cerro Solo (SCS), Arroyo Cebollín (SCE), Arroyo Pericos (SPE), Arroyo El Desecho (CED), Arroyo El Río (CER), Arroyo El Río 2 (CER2), Arroyo San Juan del Negro (CSJN) and Arroyo Agua Blanca (CAB). Additionally, trout from aquaculture facilities are included. El Barro Aquaculture Farm (AQBA), San Miguel Aquaculture Farm (AQSM), Mexican golden trout generated in laboratory (REP), and hybrids from Mexican golden trout and rainbow trout, produced in laboratory (H). The *F_ST_* coefficients among simple sites at the same basin are shown in colors: Río Fuerte in yellow, Río Sinaloa in green, Río Culiacán in red, Lab trout in orange and aquaculture trout in blue.

**Table S3.2.**
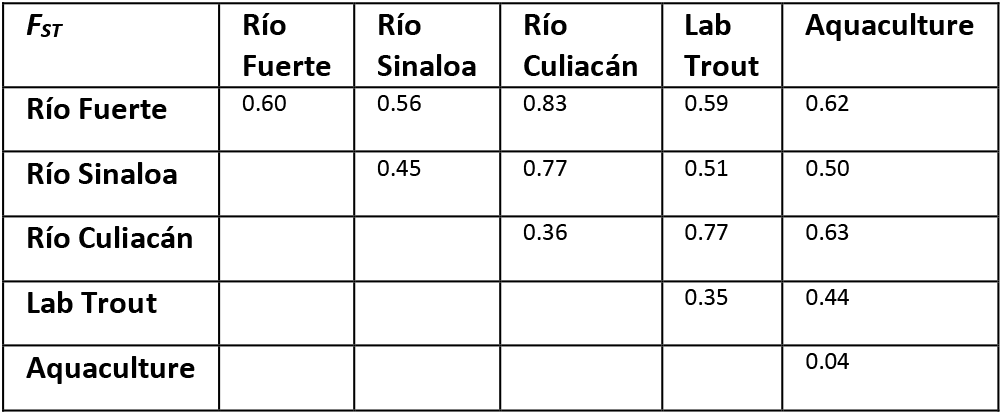
Avergae *F_ST_* coefficients within and among basins, lab trout and aquaculture farms.

## Online Resource 4

Ancestry coefficients obtained in faststructure for different K values (6, 7 and 8).

**Figure S4.1.**
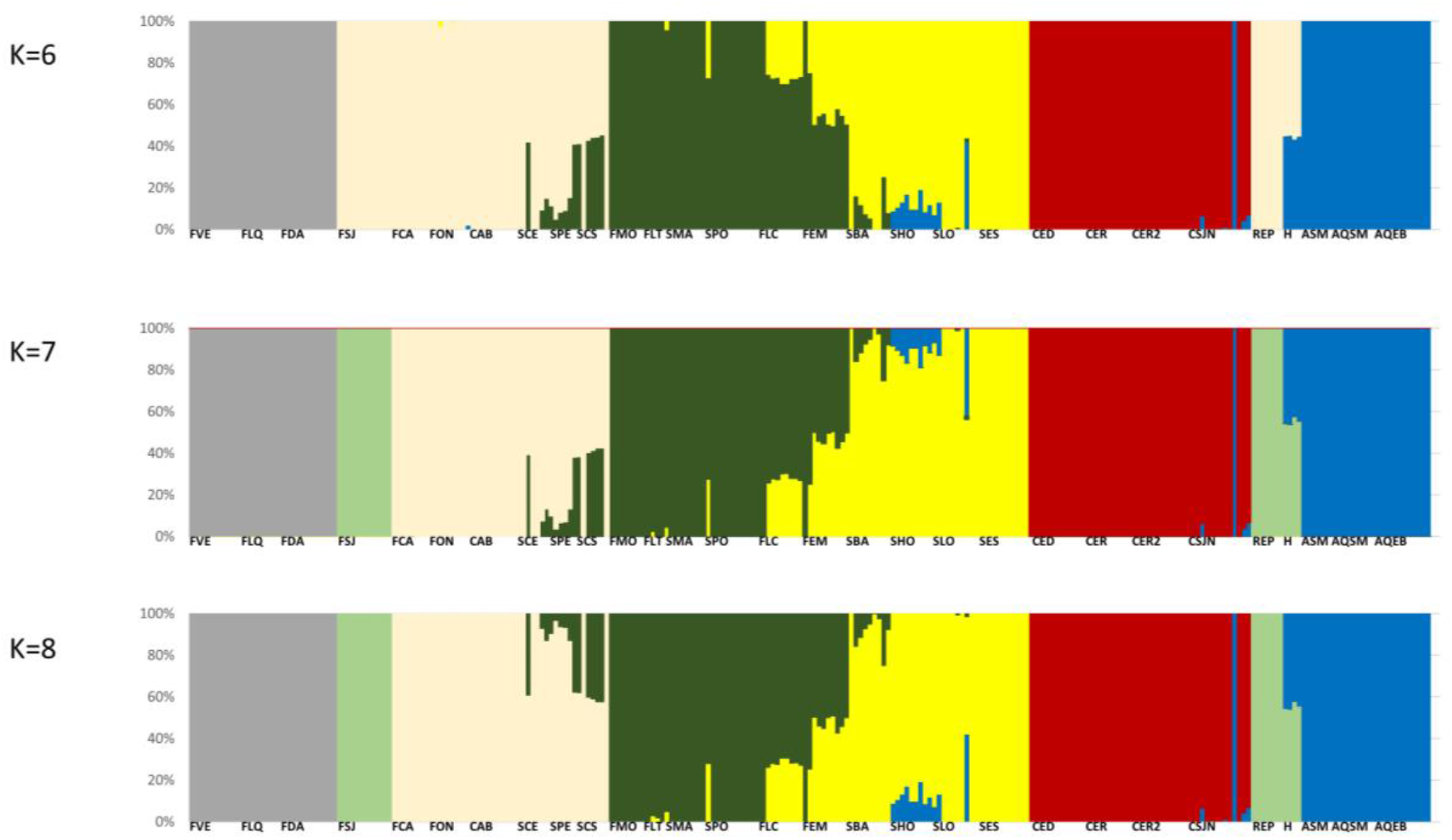
Ancestry coefficients obtained in faststructure for different K values (6, 7 and 8). Each color represents a different genetic cluster, the genome of each individual is represented by a vertical line and each sample site is represented by codes, including the initial of the river basin (F. Río Fuerte; S. Río Sinaloa; C. Río Culiacán) and an acronym of the locality Spanish name: Arroyo del Agua (FDA), Arroyo El Manzano (FEM), Arroyo Las Cuevas (FLC), Arroyo La Quebrada (FLQ), Arroyo Las Truchas (FLT), Arroyo Momorita (FMO), Arroyo San José (FSJ), Arroyo Caleras (FCA), Arroyo La Onza (FON), Arroyo Río Verde (FVE), Arroyo Baluarte (SBA), Arroyo El Salto (SES), Arroyo Hondo (SHO), Arroyo La Osera (SLO), Arroyo Macheras (SMA), Arroyo El Potrero (SPO), Arroyo San Miguel (SSM), Arroyo Cerro Solo (SCS), Arroyo Cebollín (SCE), Arroyo Pericos (SPE), Arroyo El Desecho (CED), Arroyo El Río (CER), Arroyo El Río 2 (CER2), Arroyo San Juan del Negro (CSJN) and Arroyo Agua Blanca (CAB). Additionally, trout from aquaculture facilities are included. El Barro Aquaculture Farm (AQBA), San Miguel Aquaculture Farm (AQSM), Mexican golden trout generated in laboratory (REP), and hybrids from Mexican golden trout and rainbow trout, produced in laboratory (H).

## Online Resource 5

Ancestry coefficients obtained in faststructure for Dataset B with native trout without significant introgression.

**Figure S5.1.**
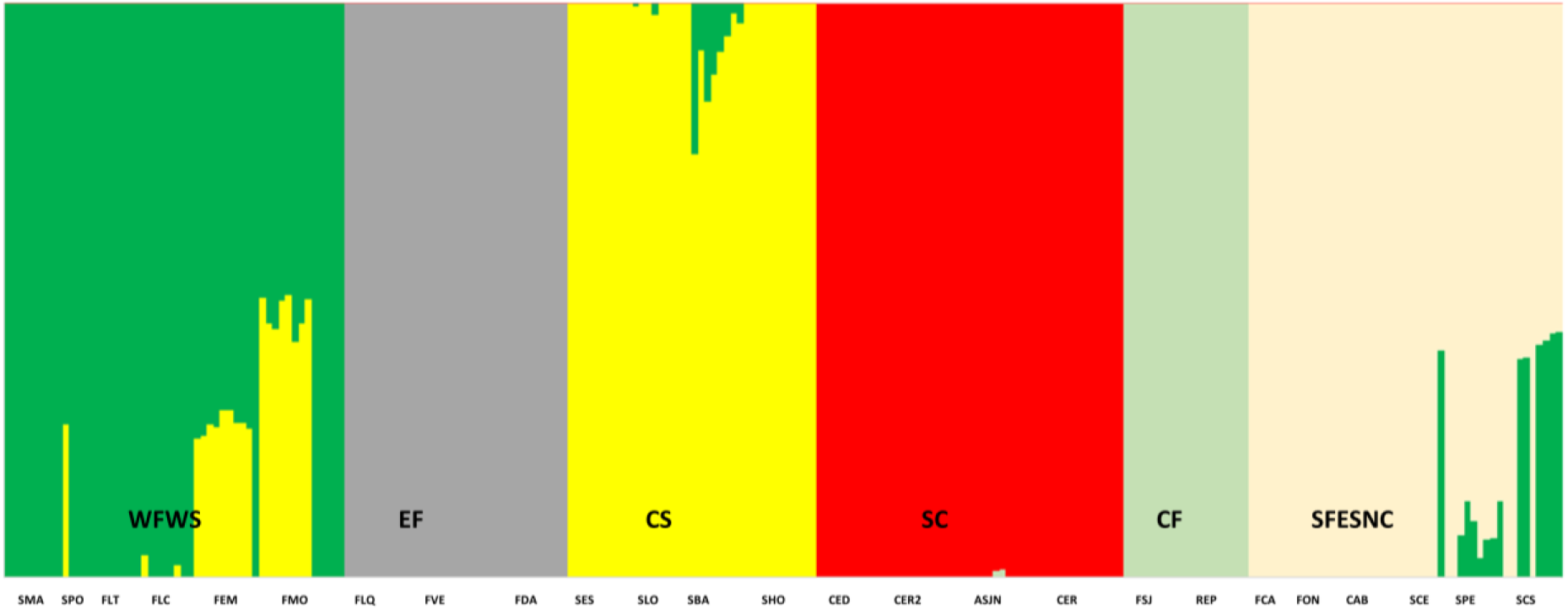
Ancestry coefficients obtained in faststructure for Dataset B with native trout without significant introgression. Each color represents a different genetic cluster: Eastern Fuerte (EF); Central Fuerte (CF); Southern Fuerte, Eastern Sinaloa and Northern Culiacán (SFESNC); Western Fuerte and Western Sinaloa (WFWS); Central Sinaloa (CS); Southern Culiacán (SC). The genome of each individual is represented by a vertical line and each sample site is represented by codes, including the initial of the river basin (F. Río Fuerte; S. Río Sinaloa; C. Río Culiacán) and an acronym of the locality Spanish name: Arroyo del Agua (FDA), Arroyo El Manzano (FEM), Arroyo Las Cuevas (FLC), Arroyo La Quebrada (FLQ), Arroyo Las Truchas (FLT), Arroyo Momorita (FMO), Arroyo San José (FSJ), Arroyo Caleras (FCA), Arroyo La Onza (FON), Arroyo Río Verde (FVE), Arroyo Baluarte (SBA), Arroyo El Salto (SES), Arroyo Hondo (SHO), Arroyo La Osera (SLO), Arroyo Macheras (SMA), Arroyo El Potrero (SPO), Arroyo Cerro Solo (SCS), Arroyo Cebollín (SCE), Arroyo Pericos (SPE), Arroyo El Desecho (CED), Arroyo El Río (CER), Arroyo El Río 2 (CER2), Arroyo San Juan del Negro (CSJN) and Arroyo Agua Blanca (CAB). Additionally, Mexican golden trout generated in laboratory (REP) are included.

## REFERENCES

Abadía-Cardoso A, Garza JC, Mayden RL, García-De León FJ (2015) Genetic Structure of Pacific Trout at the Extreme Southern End of Their Native Range. PLoS One 10:e0141775. doi: 10.1371/journal.pone.0141775

Ahrens CW, Rymer PD, Stow A, et al (2018) The search for loci under selection: trends, biases and progress. Mol Ecol 27:1342–1356. doi: 10.1111/mec.14549

Allendorf FW, Leary RF, Spruell P, Wenburg JK (2001) The problems with hybrids: Setting conservation guidelines. Trends Ecol. Evol. 16:613–622

Amish SJ, Ali O, Peacock M, et al (2019) Assessing thermal adaptation using family‐based association and F_ST_ ‐outlier tests in a threatened trout. Mol Ecol mec.15100. doi: 10.1111/mec.15100

Andrews S (2010) FastQC: a quality control tool for high throughput sequence data

Baguette M, Blanchet S, Legrand D, et al (2013) Individual dispersal, landscape connectivity and ecological networks. Biol Rev 88:310–326. doi: 10.1111/brv.12000

Bahls P (1992) The status of fish populations and management of high mountain lakes in the western United States. Northwest Sci 66:183–193. doi: 10.1016/0006-3207(94)90036-1

Banks SC, Cary GJ, Smith AL, et al (2013) How does ecological disturbance influence genetic diversity? Trends Ecol Evol 28:670–679

Behnke RL, Tomelleri JR, McGuane T (2002) Trout and Salmon of North America. Free Press, Chicago

Berthelot C, Brunet F, Chalopin D, et al (2014) The rainbow trout genome provides novel insights into evolution after whole-genome duplication in vertebrates. Nat Commun 5:3657. doi: 10.1038/ncomms4657

Binder TR, Thompson HT, Muir AM, et al (2015) New insight into the spawning behavior of lake trout, Salvelinus namaycush, from a recovering population in the Laurentian Great Lakes. Environ Biol Fishes 98:173–181. doi: 10.1007/s10641-014-0247-6

Bourret V, Dionne M, Kent MP, et al (2013) Landscape genomics in Atlantic salmon (Salmo salar): searching for gene-environment interactions driving local adaptation. Evolution 67:3469–87. doi: 10.1111/evo.12139

Brauer CJ, Hammer MP, Beheregaray LB (2016) Riverscape genomics of a threatened fish across a hydroclimatically heterogeneous river basin. Mol Ecol 25:5093–5113. doi: 10.1111/mec.13830

Brauer CJ, Unmack PJ, Smith S, et al (2018) On the roles of landscape heterogeneity and environmental variation in determining population genomic structure in a dendritic system. Mol Ecol 27:3484–3497. doi: 10.1111/mec.14808

Buchinger TJ, Marsden | J Ellen, Binder TR, et al (2017) Temporal constraints on the potential role of fry odors as cues of past reproductive success for spawning lake trout. 10196 | Ecol Evol 7:10196–10206. doi: 10.1002/ece3.3546

Camarena-Rosales F, Ruiz-Campos G, De La Rosa-Vélez J, et al (2008) Mitochondrial haplotype variation in wild trout populations (Teleostei: Salmonidae) from northwestern Mexico. Rev Fish Biol Fish 18:33–45. doi: 10.1007/s11160-007-9060-z

Carim KJ, Eby LA, Barfoot CA, Boyer MC (2016) Consistent loss of genetic diversity in isolated cutthroat trout populations independent of habitat size and quality. Conserv Genet 17:1363–1376. doi: 10.1007/s10592-016-0867-9

Catchen JM (2013) Stacks: an analysis tool set for population genomics. Mol Ecol 22:3124–3140. doi: 10.1111/mec.12354.Stacks

Catchen JM, Amores A, Hohenlohe P, et al (2011) Stacks: Building and Genotyping Loci De Novo From Short-Read Sequences. Genes|Genomes|Genetics 1:171–182. doi: 10.1534/g3.111.000240

Channell R, Lomolino M V. (2000) Dynamic biogeography and conservation of endangered species. Nature 403:84–86. doi: 10.1038/47487

Chaput-Bardy A, Lemaire C, Picard D, Secondi J (2008) In-stream and overland dispersal across a river network influences gene flow in a freshwater insect, Calopteryx splendens. Mol Ecol 17:3496–3505. doi: 10.1111/j.1365-294X.2008.03856.x

Crandall KA, Bininda-Emonds ORP, Mace GM, Wayne RK (2000) Considering evolutionary processes in conservation biology. Trends Ecol Evol 15:290–295. doi: 10.1016/S0169-5347(00)01876-0

Crockett EL (1998) Cholesterol Function in Plasma Membranes from Ectotherms: Membrane-Specific Roles in Adaptation to Temperature. Am Zool 38:291–304. doi: 10.1093/icb/38.2.291

Crozier LG, Hutchings JA (2014) Plastic and evolutionary responses to climate change in fish. Evol Appl 7:68–87. doi: 10.1111/eva.12135

Cushman SA, Landguth EL (2010) Spurious correlations and inference in landscape genetics. Mol Ecol 19:3592–3602. doi: 10.1111/j.1365-294X.2010.04656.x

Dalongeville A, Benestan L, Mouillot D, et al (2018) Combining six genome scan methods to detect candidate genes to salinity in the Mediterranean striped red mullet (Mullus surmuletus). BMC Genomics 19:217. doi: 10.1186/s12864-018-4579-z

Danecek P, Auton A, Abecasis G, et al (2011) The variant call format and VCFtools. Bioinformatics 27:2156–2158. doi: 10.1093/bioinformatics/btr330

Davis CD, Epps CW, Flitcroft RL, Banks MA (2018) Refining and defining riverscape genetics: How rivers influence population genetic structure. Wiley Interdiscip Rev Water 5:e1269. doi: 10.1002/wat2.1269

de Lafontaine G, Napier JD, Petit RJ, Hu FS (2018) Invoking adaptation to decipher the genetic legacy of past climate change. Ecology 99:1530–1546. doi: 10.1002/ecy.2382

Dijkstra EW (1959) A Note on Two Problems in Connexion with Graphs. Numer Math 1:269–27

Do C, Waples RS, Peel D, et al (2014) NeEstimator v2: Re-implementation of software for the estimation of contemporary effective population size (Ne) from genetic data. Mol Ecol Resour 14:209–214. doi: 10.1111/1755-0998.12157

Eckert CG, Samis KE, Lougheed SC (2008) Genetic variation across species’ geographical ranges: The central-marginal hypothesis and beyond. Mol Ecol 17:1170–1188. doi: 10.1111/j.1365-294X.2007.03659.x

Elshire RJ, Glaubitz JC, Sun Q, et al (2011) A robust, simple genotyping-by-sequencing (GBS) approach for high diversity species. PLoS One 6:1–10. doi: 10.1371/journal.pone.0019379

Escalante MA, García-De-León FJ, Dillman CB, et al (2014) Genetic introgression of cultured rainbow trout in the Mexican native trout complex. Conserv Genet 15:1063–1071. doi: 10.1007/s10592-014-0599-7

Escalante MA, García-De León FJ, Dillman CB, et al (2016) Introgresión genética de la trucha arcoíris exótica en poblaciones de trucha dorada mexicana. In: Ruiz-Luna A, García-De León FJ (eds) La Trucha Dorada Mexicana. CIAD-CIBNOR, Mexico, pp 125–136

Escalante MA, García-De León FJ, Ruiz-Luna A, et al (2018) The interplay of riverscape features and exotic introgression on the genetic structure of the Mexican golden trout (Oncorhynchus chrysogaster), a simulation approach. J Biogeogr 45:1500–1514. doi: 10.1111/jbi.13246

Fan H, Hu Y, Wu Q, et al (2018) Conservation genetics and genomics of threatened vertebrates in China. J Genet Genomics 45:593–601. doi: 10.1016/J.JGG.2018.09.005

Faulks LK, Kerezsy A, Unmack PJ, et al (2017) Going, going, gone? Loss of genetic diversity in two critically endangered Australian freshwater fishes, Scaturiginichthys vermeilipinnis and Chlamydogobius squamigenus, from Great Artesian Basin springs at Edgbaston, Queensland, Australia. Aquat Conserv Mar Freshw Ecosyst 27:39–50. doi: 10.1002/aqc.2684

Fausch KD (2007) Introduction, establishment and effects of non-native salmonids: Considering the risk of rainbow trout invasion in the United Kingdom. J Fish Biol 71:1–32. doi: 10.1111/j.1095-8649.2007.01682.x

Flanagan SP, Forester BR, Latch EK, et al (2018) Guidelines for planning genomic assessment and monitoring of locally adaptive variation to inform species conservation. Evol Appl 11:1035–1052. doi: 10.1111/eva.12569

Frichot E, François O (2015) LEA: An R package for landscape and ecological association studies. Methods Ecol Evol 6:925–929. doi: 10.1111/2041-210X.12382

Frichot E, Schoville SD, Bouchard G, François O (2013) Testing for associations between loci and environmental gradients using latent factor mixed models. Mol Biol Evol 30:1687–1699. doi: 10.1093/molbev/mst063

Gagnaire P-A, Gaggiotti OE (2016) Detecting polygenic selection in marine populations by combining population genomics and quantitative genetics approaches. Curr Zool 1–14. doi: 10.1093/cz/zow088

García-De León FJ, Dillman CB, De Los Santos Camarillo AB, et al (2020) First steps towards the identification of evolutionarily significant units in Mexican native trout: An assessment of microsatellite variation. Environ Biol Fishes. doi: 10.1007/s10641-020-00979-4

García-De León FJ, Getino-Mamet LN, Rodríguez-Jaramillo C, et al (2016) Dimorfismo sexual y periodo reproductivo de la trucha dorada mexicana, Oncorhynchus chrysogaster en los ríos Fuerte, Sinaloa y Culiacán. In: Ruíz-Luna A, García-De León FJ (eds) La Trucha Dorada Mexicana. CIAD-CIBNOR, pp 53–72

Grummer JA, Beheregaray LB, Bernatchez L, et al (2019) Aquatic Landscape Genomics and Environmental Effects on Genetic Variation. Trends Ecol Evol. doi: 10.1016/j.tree.2019.02.013

Gunnell K, Tada MK, Hawthorne FA, et al (2008) Geographic patterns of introgressive hybridization between native Yellowstone cutthroat trout (Oncorhynchus clarkii bouvieri) and introduced rainbow trout (O. mykiss) in the South Fork of the Snake River watershed, Idaho. Conserv Genet 9:49–64. doi: 10.1007/s10592-007-9302-6

Günther T, Coop G (2013) Robust identification of local adaptation from allele frequencies. Genetics 195:205–220. doi: 10.1534/genetics.113.152462

Hale MC, Xu P, Scardina J, et al (2011) Differential gene expression in male and female rainbow trout embryos prior to the onset of gross morphological differentiation of the gonads. BMC Genomics 12:404. doi: 10.1186/1471-2164-12-404

Hand BK, Muhlfeld CC, Wade AA, et al (2016) Climate variables explain neutral and adaptive variation within salmonid metapopulations: the importance of replication in landscape genetics. Mol Ecol 25:689–705. doi: 10.1111/mec.13517

Hecht BC, Matala AP, Hess JE, Narum SR (2015) Environmental adaptation in Chinook salmon (Oncorhynchus tshawytscha) throughout their North American range. Mol Ecol 24:5573–5595. doi: 10.1111/mec.13409

Heggberget TG, Johnsen BO, Hindar K, et al (1993) Interactions between wild and cultured Atlantic salmon: a review of the Norwegian experience. Fish Res 18:123–146. doi: 10.1016/0165-7836(93)90044-8

Hendricks S, Anderson EC, Antao T, et al (2018) Recent advances in conservation and population genomics data analysis. Evol Appl 11:1197–1211. doi: 10.1111/eva.12659

Hendrickson DA, Neely DA, Mayden RL, et al (2006) Conservation of Mexican native trout and the discovery, status. Protection and recovery of the Conchos trout, the first native. Stud North Am desert fishes Honor EP Pist Conserv Univ Autónoma Nuevo León, Monterrey 162–201

Hendrickson DA, Pérez HE, Findley LT, et al (2002) Mexican native trouts: A review of their history and current systematic and conservation status. Rev Fish Biol Fish 12:273–316. doi: 10.1023/A:1025062415188

Hewitt G (2000) The genetic legacy of the Quaternary ice ages. Nature 405:907–913. doi: 10.1038/35016000

Hoban S, Kelley JL, Lotterhos KE, et al (2016) Finding the Genomic Basis of Local Adaptation: Pitfalls, Practical Solutions, and Future Directions. Am Nat 188:379–97. doi: 10.1086/688018

Höglund J (2009) Evolutionary conservation genetics. Oxford University Press

Hogstrand C, Wilson RW, Polgar D, Wood CM (1994) Effects of zinc on the kinetics of branchial calcium uptake in freshwater rainbow trout during adaptation to waterborne zinc. J Exp Biol 186:55–73

Hunter ME, Hoban SM, Bruford MW, et al (2018) Next-generation conservation genetics and biodiversity monitoring. Evol Appl 11:1029–1034. doi: 10.1111/eva.12661

Johnson NS, Higgs D, Binder TR, et al (2018) Evidence of sound production by spawning lake trout (*Salvelinus namaycush*) in lakes Huron and Champlain. Can J Fish Aquat Sci 75:429–438. doi: 10.1139/cjfas-2016-0511

Jombart T (2008) Adegenet: A R package for the multivariate analysis of genetic markers. Bioinformatics 24:1403–1405. doi: 10.1093/bioinformatics/btn129

Kanno Y, Vokoun JC, Letcher BH (2011) Fine-scale population structure and riverscape genetics of brook trout (Salvelinus fontinalis) distributed continuously along headwater channel networks. Mol Ecol 20:3711–3729. doi: 10.1111/j.1365-294X.2011.05210.x

Kleinman-Ruiz D, Martínez-Cruz B, Soriano L, et al (2017) Novel efficient genome-wide SNP panels for the conservation of the highly endangered Iberian lynx. BMC Genomics 18:556. doi: 10.1186/s12864-017-3946-5

Kokko H, Chaturvedi A, Croll D, et al (2017) Can Evolution Supply What Ecology Demands? Trends Ecol Evol 32:187–197. doi: 10.1016/j.tree.2016.12.005

Koskinen MT, Haugen TO, Primmer CR (2002) Contemporary fisherian life-history evolution in small salmonid populations. Nature 419:826–830. doi: 10.1038/nature01029

Landguth EL, Bearlin A, Day CC, Dunham J (2016) CDMetaPOP: an individual-based, eco-evolutionary model for spatially explicit simulation of landscape demogenetics. Methods Ecol Evol 8:4–11. doi: 10.1111/2041-210X.12608

Landguth EL, Fedy BC, Oyler-Mccance SJ, et al (2012) Effects of sample size, number of markers, and allelic richness on the detection of spatial genetic pattern. Mol Ecol Resour 12:276–284. doi: 10.1111/j.1755-0998.2011.03077.x

Lawler JJ, Tear TH, Pyke C, et al (2010) Resource management in a changing and uncertain climate. Front Ecol Environ 8:35–43. doi: 10.1890/070146

Le Pichon CLE, Gorges G, Boët P, et al (2006) A spatially explicit resource-based approach for managing stream fishes in riverscapes. Environ Manage 37:322–335. doi: 10.1007/s00267-005-0027-3

Leitwein M, Garza JC, Pearse DE (2017) Ancestry and adaptive evolution of anadromous, resident, and adfluvial rainbow trout (Oncorhynchus mykiss) in the San Francisco bay area: application of adaptive genomic variation to conservation in a highly impacted landscape. Evol Appl 10:56–67. doi: 10.1111/eva.12416

Linløkken AN, Haugen TO, Mathew PK, et al (2016) Comparing estimates of number of breeders Nb based on microsatellites and single nucleotide polymorphism of three groups of brown trout (Salmo trutta L.). Fish Manag Ecol 23:152–160. doi: 10.1111/fme.12169

Luu K, Bazin E, Blum MGB (2016) *pcadapt* : an R package to perform genome scans for selection based on principal component analysis. Mol Ecol Resour. doi: 10.1111/1755-0998.12592

Manel S, Holderegger R (2013) Ten years of landscape genetics. Trends Ecol Evol 28:614–621. doi: 10.1016/j.tree.2013.05.012

Marie AD, Bernatchez L, Garant D, Taylor E (2012) Environmental factors correlate with hybridization in stocked brook charr (*Salvelinus fontinalis*). Can J Fish Aquat Sci 69:884–893. doi: 10.1139/f2012-027

Martin M (2011) Sequencing Reads. EMBnet.journal 17:10–12

Meirmans PG, Van Tienderen PH (2004) GENOTYPE and GENODIVE: Two programs for the analysis of genetic diversity of asexual organisms. Mol Ecol Notes 4:792–794. doi: 10.1111/j.1471-8286.2004.00770.x

Milanesi P, Holderegger R, Caniglia R, et al (2017) Expert-based versus habitat-suitability models to develop resistance surfaces in landscape genetics. Oecologia 183:67–79. doi: 10.1007/s00442-016-3751-x

Milano I, Babbucci M, Cariani A, et al (2014) Outlier SNP markers reveal fine-scale genetic structuring across European hake populations (Merluccius merluccius). Mol Ecol 23:118–135. doi: 10.1111/mec.12568

Milián-García Y, Ramos-Targarona R, Pérez-Fleitas E, et al (2015) Genetic evidence of hybridization between the critically endangered Cuban crocodile and the American crocodile: implications for population history and in situ/ex situ conservation. Heredity (Edinb) 114:272–280. doi: 10.1038/hdy.2014.96

Miller RR, Williams JD, Williams JE (1989) Extinctions of North American Fishes During the Past Century. Fisheries 14:22–38. doi: 10.2307/1104578

Morita K, Yamamoto S (2002) Effects of habitat fragmentation by damming on the persistence of a stream - dwelling charr. Conserv Biol 16:1318–1323. doi: 10.1046/j.1523-1739.2002.01476.x

Muhlfeld CC, Kalinowski ST, McMahon TE, et al (2009) Hybridization rapidly reduces fitness of a native trout in the wild. Biol Lett

Muhlfeld CC, Kovach RP, Al-Chokhachy R, et al (2017) Legacy introductions and climatic variation explain spatiotemporal patterns of invasive hybridization in a native trout. Glob Chang Biol 23:4663–4674. doi: 10.1111/gcb.13681

Narum SR, Zendt JS, Graves D, Sharp WR (2008) Influence of landscape on resident and anadromous life history types of *Oncorhynchus mykiss*. Can J Fish Aquat Sci 65:1013–1023. doi: 10.1139/F08-025

Needham PR, Gard R (1964) A New Trout from Central Mexico: Salmo chrysogaster, the Mexican Golden Trout. Copeia 1964:169. doi: 10.2307/1440847

Nielsen JL, Sage GK (2001) Microsatellite analyses of the trout of northwest Mexico. Genetica 111:269–278. doi: 10.1023/A:1013777701213

Nomura T (2008) Estimation of effective number of breeders from molecular coancestry of single cohort sample. Evol Appl 1:462–474. doi: 10.1111/j.1752-4571.2008.00015.x

Olsen MT, Islas V, Graves JA, et al (2017) Genetic population structure of harbour seals in the United Kingdom and neighbouring waters. doi: 10.1002/aqc.2760

Parmesan C (2006) Ecological and Evolutionary Responses to Recent Climate Change. Annu Rev Ecol Evol Syst 37:637–669. doi: 10.1146/annurev.ecolsys.37.091305.110100

Pearse DE (2016) Saving the spandrels? Adaptive genomic variation in conservation and fisheries management. J Fish Biol 89:2697–2716. doi: 10.1111/jfb.13168

Pearse DE, Campbell MA (2018) Ancestry and Adaptation of Rainbow Trout in Yosemite National Park. Fisheries 43:472–484. doi: 10.1002/fsh.10136

Penaluna BE, Abadía-Cardoso A, Dunham JB, et al (2016) Conservation of Native Pacific Trout Diversity in Western North America. Fisheries 41:286–300. doi: 10.1080/03632415.2016.1175888

Perreault-Payette A, Muir AM, Goetz F, et al (2017) Investigating the extent of parallelism in morphological and genomic divergence among lake trout ecotypes in Lake Superior. Mol Ecol 26:1477–1497. doi: 10.1111/mec.14018

Perrier C, Ferchaud A-L, Sirois P, et al (2017) Do genetic drift and accumulation of deleterious mutations preclude adaptation? Empirical investigation using RADseq in a northern lacustrine fish. Mol Ecol 26:6317–6335. doi: 10.1111/mec.14361

Perrier C, Guyomard R, Bagliniere J-L, et al (2013) Changes in the genetic structure of Atlantic salmon populations over four decades reveal substantial impacts of stocking and potential resiliency. Ecol Evol 3:2334–2349. doi: 10.1002/ece3.629

Perrier C, Guyomard R, Bagliniere JL, Evanno G (2011) Determinants of hierarchical genetic structure in Atlantic salmon populations: Environmental factors vs. anthropogenic influences. Mol Ecol 20:4231–4245. doi: 10.1111/j.1365-294X.2011.05266.x

Rahel FJ, Bierwagen B, Taniguchi Y (2008) Managing aquatic species of conservation concern in the face of climate change and invasive species. Conserv Biol 22:551–561. doi: 10.1111/j.1523-1739.2008.00953.x

Raj A, Stephens M, Pritchard JK (2014) FastSTRUCTURE: Variational inference of population structure in large SNP data sets. Genetics 197:573–589. doi: 10.1534/genetics.114.164350

Razgour O, Forester B, Taggart JB, et al (2019) Considering adaptive genetic variation in climate change vulnerability assessment reduces species range loss projections. Proc Natl Acad Sci 116:10418–10423. doi: 10.1073/pnas.1820663116

Richardson JL, Brady SP, Wang IJ, Spear SF (2016) Navigating the pitfalls and promise of landscape genetics. Mol Ecol 25:849–863. doi: 10.1111/mec.13527

Rieman BE, Allendorf FW (2001) Effective population size and genetic conservation criteria for bull trout. North Am J Fish Manag 21:756–764. doi: 10.1577/1548-8675(2001)021<0756:EPSAGC>2.0.CO;2

Riginos C, Liggins L (2013) Seascape Genetics: Populations, Individuals, and Genes Marooned and Adrift. Geogr Compass 7:197–216. doi: 10.1111/gec3.12032

Rosenberg MS, Anderson CD (2011) PASSaGE: Pattern Analysis, Spatial Statistics and Geographic Exegesis. Version 2. Methods Ecol Evol 2:229–232. doi: 10.1111/j.2041-210X.2010.00081.x

Ruiz-Luna A, Hernández-Guzmán R, García-De León FJ, Ramírez-Huerta AL (2017) Potential distribution of endangered Mexican golden trout (Oncorhynchus chrysogaster) in the Rio Sinaloa and Rio Culiacan basins (Sierra Madre Occidental) based on landscape characterization and species distribution models. Environ Biol Fishes 1–28. doi: 10.1007/s10641-017-0624-z

Ruzzante DE, McCracken GR, Parmelee S, et al (2016) Effective number of breeders, effective population size and their relationship with census size in an iteroparous species, Salvelinus fontinalis. Proc R Soc B 283:20152601. doi: 10.1098/rspb.2015.2601

Saitou N, Nei M (1987) The neighbour-joining method: a new method for reconstructing phylogenetic trees. Mol Biol Evo 4:406–425. doi: citeulike-article-id:93683

Salem M, Kenney PB, Rexroad CE, Yao J (2010) Proteomic signature of muscle atrophy in rainbow trout. J Proteomics 73:778–789. doi: 10.1016/j.jprot.2009.10.014

Schmidt DJ, Espinoza T, Connell M, Hughes JM (2017) Conservation genetics of the Mary River turtle (Elusor macrurus) in natural and captive populations. doi: 10.1002/aqc.2851

Splendiani A, Ruggeri P, Giovannotti M, Caputo Barucchi V (2013) Role of environmental factors in the spread of domestic trout in Mediterranean streams. Freshw Biol 58:2089–2101. doi: 10.1111/fwb.12193

Taggart JB, Bron JE, Martin SAM, et al (2008) A description of the origins, design and performance of the TRAITS-SGP Atlantic salmon Salmo salar L. cDNA microarray. J Fish Biol 72:2071–2094. doi: 10.1111/j.1095-8649.2008.01876.x

Taiyun Wei M (2017) Title Visualization of a Correlation Matrix. Statistician 56:316–324

Tamura K, Nei M (1993) Estimation of the number of nucleotide substitutions in the control region of mitochondrial DNA in humans and chimpanzees. Mol Biol Evol 10:512–26. doi: 10.1093/molbev/msl149

Team R. Core (2018) R: A language and environment for statistical computing. 0:201. doi: 10.1108/eb003648

Tikochinski Y, Bradshaw P, Mastrogiacomo A, et al (2018) Mitochondrial DNA short tandem repeats unveil hidden population structuring and migration routes of an endangered marine turtle. Aquat Conserv Mar Freshw Ecosyst. doi: 10.1002/aqc.2908

Torres-Florez JP, Johnson WE, Nery MF, et al (2017) The coming of age of conservation genetics in Latin America: what has been achieved and what needs to be done. Conserv Genet 19:1–15. doi: 10.1007/s10592-017-1006-y

Torterotot J-B, Perrier C, Bergeron NE, Bernatchez L (2014) Influence of forest road culverts and waterfalls on the fine-scale distribution of brook trout genetic diversity in a boreal watershed. Trans Am Fish Soc 143:1577–1591. doi: 10.1080/00028487.2014.952449

van Etten J (2012) gdistance: Distances and routes on geographical grids. R package version 1.1--4. Available at CRANR-projectorg/package=gdistance

Vera M, Martinez P, Bouza C (2017) Stocking impact, population structure and conservation of wild brown trout populations in inner Galicia (NW Spain), an unstable hydrologic region. Aquat Conserv Mar Freshw Ecosyst. doi: 10.1002/aqc.2856

Weigel DE, Peterson JT, Spruell P (2003) Introgressive Hybridization between Native Cutthroat Trout and Introduced Rainbow Trout. Ecol. Appl. 13:38–50

Wenne R, Bernaś R, Poćwierz-Kotus A, et al (2016) Recent genetic changes in enhanced populations of sea trout (Salmo trutta m. trutta) in the southern Baltic rivers revealed with SNP analysis. Aquat Living Resour 103:29. doi: 10.1051/alr/2016012

Winans GA, Allen MB, Baker J, et al (2018) Dam trout: Genetic variability in Oncorhynchus mykiss above and below barriers in three Columbia River systems prior to restoring migrational access. PLoS One 13:e0197571. doi: 10.1371/journal.pone.0197571

## LITERATURE

ESRI. (2013). ArcGIS, version 10.2 for Desktop. ESRI, Redlands, California.

Escalante, M. A., García-De León, F. J., Ruiz-Luna, A., Landguth, E., & Manel, S. (2018). The interplay of riverscape features and exotic introgression on the genetic structure of the Mexican golden trout (*Oncorhynchus chrysogaster*), a simulation approach. Journal of Biogeography, 45(7), 1500–1514. https://doi.org/10.1111/jbi.13246

van Etten, J. (2012). gdistance: Distances and routes on geographical grids. R package version 1.1--4. Available at CRAN.R-Project.Org/Package=Gdistance.

